# Astrocyte-driven small vessel disease is an early, amyloid-independent feature of PSEN1 E280A familial Alzheimer’s disease

**DOI:** 10.64898/2026.07.17.739026

**Authors:** Jessica Lisa Villalba-Moreno, Yassir El-Amri, Keun-Young Kim, Nelson David Villalba-Moreno, Mohsin Shafiq, Carolina Ortiz-Cordero, Shiwei Wang, J Alejandro Ossa, Isaura Suarez-Uribe, Duvan Cardona-Madrigal, Andres Villegas, Markus Glatzel, Susanne Krasemann, Rafael Posada-Duque, Laura L Kiessling, Francisco Lopera, Joseph Arboleda-Velasquez, Rajesh Kalaria, Mark Ellisman, Diego Sepulveda-Falla

## Abstract

Cerebral Small vessel disease (cSVD) is a prevalent feature of Alzheimer’s disease (AD) pathology. Whether this pathology is a late consequence of amyloid and tau accumulation or an early, direct effect of PSEN1 dysfunction has remained unresolved. We found that it is more severe in familial AD (FAD) caused by E280A mutation in presenilin 1 (PSEN1). These cases present with a distinctive proteomic signature, associated with pathological features, more dysregulated in the occipital cortex (OC) compared to the frontal cortex (FC), and characterized by multiple dysregulated proteins involved in extracellular matrix (ECM) and RNA-associated processes. This proteomic fingerprint was associated with abnormal collagen build up, ECM disorganization, and signatures of aberrant angiogenesis. Six months old transgenic knock-in mice homozygous for Psen1 E280A mutation (PSEN1Ki) also showed a similar phenotype with microvascular tortuosity and proteomic changes. Critically, these mice develop neither Aβ plaques nor tau tangles, indicating that the shared microvascular and RNA-associated changes are direct consequences of PSEN1 dysfunction rather than downstream effects of amyloid pathology. Remarkably, dysregulated RNA-associated protein networks overlapped between FAD and PSEN1Ki mice. Cerebral microvessels microstructure in PSEN1Ki mice at two months and six months showed abnormal astrocytic end-feet with lamellar deposits implicating blood-brain barrier damage. Finally, single nuclei transcriptomic analysis of AD patients and controls showed similar abnormal astrocytes in both sporadic and familial variants, but FAD astrocytes expressed dysregulated genes identified in the proteomic analyses, such as GLUL, APOE, and CLU. Our findings suggest that cSVD is an early pathological event in PSEN1 FAD and that is driven by abnormal RNA-associated processes and astrocytic dysfunction.

## Introduction

The presence of amyloid-beta (Aβ) plaques and hyperphosphorylated tau deposits are required for an Alzheimer’s Disease (AD) diagnosis. AD pathology also includes Aβ deposits in the walls of small vessels, known as cerebral amyloid angiopathy (CAA). Several pathophysiological processes contribute to AD, including the accumulation of misfolded proteins (Aβ and tau), neuroinflammation, and cerebral vascular dysfunction^1^.

Familial Alzheimer’s disease (FAD). is caused by mutations in the amyloid precursor protein (APP) or presenilin (PSEN1 or 2) genes, with an earlier onset, commonly in the fourth decade of life. FAD brains exhibit more Aβ deposits and are more prone to CAA due to increased Aβ production^2^.

The p.Glu280Ala (E280A) mutation in PSEN1 affects the largest population in the world suffering from FAD, with over 1000 confirmed carriers in the Colombian kindred. E280A carriers exhibit memory and language impairments and behavioral changes^3^. This mutation leads to severe brain atrophy, significant amyloid-beta (Aβ) pathology, hyperphosphorylated tau pathology, and cerebellar damage^4,5^. E280A carriers often have an increased prevalence of CAA compared to SAD^6^. E280A carriers also exhibit a shift towards Aβ peptides 1-38, 1-42, and 1-43^7^, and often present comorbidities like cerebrovascular disease and Lewy body dementia (LBD)^6^.

Vascular pathology primarily in form of cSVD is increasingly recognized as a major contributing factor to AD. FAD mutation carriers show greater white matter degeneration and white matter hyperintensities (WMH) indicated to be of vascular origin^8^. cSVD describes pathological changes in small end arteries, arterioles, venules, and brain capillaries, leading to reduced or interrupted organ perfusion. Major risk factors include high blood pressure, diabetes mellitus, hyperlipidemia, smoking, and cardiovascular disease. SVD can cause vascular cognitive impairment and dementia (VCID) when the brain is affected. It is the leading cause of vascular dementia (VaD) and accounts for 20% of all strokes. Early cSVD changes can include arteriolosclerosis and CAA, leading to lacunar infarcts, microinfarcts, and microbleeds. Microaneurysms and vascular resistance changes can precede amyloid deposition. Post-mortem studies have described significant vascular pathology to be present in >80% in AD patients compared to healthy individuals^8^.

We previously identified cSVD not associated with CAA or general Aβ pathology, in E280A FAD cases. Specifically, we identified increased perivascular space (PVS) and mural damage^9^. Here, we used proteomics to identify the underlying causes of the different types of cSVD pathology found in E280A cSVD. We found that besides distinct proteomic signature associated with CAA in both SAD and FAD cases, functional proteomic modules are specifically associated with CAA severity, PVS dilation and FAD. Among these functional modules, RNA regulation and processing, as well as vascular ECM was identified in E280A FAD. While we previously described Aβ-independent small vessel pathology in this population, the molecular drivers and the timing of its onset remained unknown. Here we ask whether this pathology arises directly from PSEN1 dysfunction and whether it precedes amyloid deposition. RNA-associated proteins were similarly dysregulated in a murine knock-in E280A model, which presented early vascular changes in the absence of Aβ pathology, associated with astrocytic dysfunction, similarly present in E280A FAD cases.

## Material and methods

### Human subjects

Table 1 shows the demographic details, relevant clinical and pathological features of the selected cases. FAD cases were selected based on age of onset (AoO), while SAD and healthy controls (HC) were of similar age. The PSEN1 E280A genealogy was first identified 30 years ago, and carriers were regularly followed up using the CERAD protocol, NINCDS-ADRA and DSM-IV criteria until end-stage dementia and death^10^. A total of 37 cases, (22 FAD, 10 SAD and 5 HC) were included in the current study.

**Table 1.** Demographic data of cases used in this study. Abbreviations: Age of Onset (AoO), Age of Death (AoD), Disease Duration (DD), Postmortem Interval in hours (PMI), APOE Haplotype (ApoE), Neuropathological assessment of microvascular disease (Neuropathology), Frontal cortex proteomics of isolated microvessels (FCp), Occipital cortex proteomics of isolated microvessels (OCp).

| Case | Group | Sex | AoO | AoD | DD | PMI | ApoE | Neuropathology | FC | OC |
| --- | --- | --- | --- | --- | --- | --- | --- | --- | --- | --- |
| 1 | FAD | 2 | 55 | 64 | 9 | 13 | NA | 1 | 0 | 0 |
| 2 | FAD | 1 | 57 | 60 | 3 | 4 | 33 | 1 | 0 | 1 |
| 3 | FAD | 2 | 46 | 54 | 5 | 12 | 34 | 1 | 0 | 0 |
| 4 | FAD | 2 | 45 | 50 | 5 | 7.5 | 33 | 1 | 0 | 1 |
| 5 | FAD | 1 | 44 | 52 | 8 | 4.8 | 33 | 1 | 0 | 0 |
| 6 | FAD | 2 | 55 | 64 | 9 | 3 | 33 | 1 | 0 | 1 |
| 7 | FAD | 2 | 40 | 42 | 2 | 5.5 | 33 | 1 | 1 | 1 |
| 8 | FAD | 1 | 47 | 55 | 8 | 3.3 | 33 | 1 | 0 | 1 |
| 9 | FAD | 2 | 57 | 62 | 5 | 4 | 33 | 1 | 1 | 1 |
| 10 | FAD | 2 | 45 | 48 | 3 | 4 | 33 | 1 | 1 | 1 |
| 11 | FAD | 2 | 41 | 47 | 6 | 2.3 | 33 | 1 | 1 | 1 |
| 12 | FAD | 2 | 50 | 60 | 10 | 2.8 | 33 | 1 | 0 | 0 |
| 13 | FAD | 2 | 45 | 55 | 10 | 2.5 | 23 | 1 | 0 | 0 |
| 14 | FAD | 2 | 62 | 74 | 12 | 5.5 | 33 | 1 | 0 | 1 |
| 15 | FAD | 1 | 56 | 63 | 7 | 3.75 | 33 | 1 | 1 | 1 |
| 16 | FAD | 1 | 44 | 51 | 7 | 3.9 | 33 | 1 | 1 | 1 |
| 17 | FAD | 2 | 47 | 52 | 5 | 8.5 | 33 | 1 | 0 | 0 |
| 18 | FAD | 2 | 58 | 64 | 6 | 3.5 | 24 | 1 | 0 | 1 |
| 19 | FAD | 2 | 49 | 62 | 11 | 3.5 | 33 | 1 | 0 | 0 |
| 20 | FAD | 1 | 44 | 48 | 4 | 12 | 33 | 1 | 0 | 1 |
| 21 | FAD | 1 | 45 | 53 | 8 | 7.2 | 34 | 1 | 0 | 1 |
| 22 | FAD | 1 | 55 | 61 | 6 | 7.3 | 33 | 1 | 1 | 1 |
| 23 | FAD | 2 | 49 | 57 | 8 | 7.8 | 33 | 1 | 0 | 0 |
| 24 | FAD | 1 | 58 | 64 | 6 | 4 | 33 | 1 | 1 | 1 |
| 25 | FAD | 2 | 49 | 57 | 8 | 9.2 | 33 | 1 | 0 | 0 |
| 26 | FAD | 1 | 57 | 64 | 7 | 5 | 33 | 1 | 1 | 1 |
| 27 | FAD | 1 | 42 | 47 | 5 | 2.7 | 34 | 1 | 0 | 1 |
| 28 | FAD | 1 | 47 | 62 | 15 | 4 | 34 | 1 | 0 | 0 |
| 29 | FAD | 1 | 47 | 54 | 7 | 8.3 | 33 | 1 | 0 | 0 |
| 30 | FAD | 2 | 45 | 51 | 6 | 4.7 | 33 | 1 | 1 | 1 |
| 31 | FAD | 2 | 52 | 60 | 8 | 5.1 | 33 | 1 | 0 | 1 |
| 32 | FAD | 1 | 46 | 62 | 16 | 4.3 | 33 | 1 | 0 | 0 |
| 33 | SAD | 1 | 78 | 86 | 8 | 6.3 | NA | 1 | 0 | 1 |
| 34 | SAD | 2 | 55 | 70 | 15 | 11.3 | 34 | 1 | 0 | 0 |
| 35 | SAD | 1 | 78 | 85 | 7 | 2.8 | 34 | 1 | 0 | 0 |
| 36 | SAD | 2 | 82 | 92 | 10 | 4.5 | 33 | 1 | 0 | 0 |
| 37 | SAD | 2 | 65 | 80 | 15 | 3.5 | 33 | 1 | 0 | 0 |
| 38 | SAD | 2 | 62 | 74 | 12 | 2.5 | 33 | 1 | 1 | 1 |
| 39 | SAD | 2 | 55 | 75 | 20 | 4.3 | 44 | 1 | 1 | 1 |
| 40 | SAD | 2 | 69 | 76 | 7 | 3.75 | 34 | 1 | 1 | 1 |
| 41 | SAD | 1 | 70 | 83 | 13 | 4.3 | 32 | 1 | 1 | 1 |
| 42 | SAD | 2 | 50 | 61 | 11 | 4.1 | 33 | 1 | 0 | 0 |
| 43 | Control | 2 | NA | 84 | NA | 4 | NA | NA | 1 | 1 |
| 44 | Control | 2 | NA | 67 | NA | 5 | NA | NA | 1 | 1 |
| 45 | Control | 2 | NA | 75 | NA | 4 | NA | NA | 1 | 1 |
| 46 | Control | 1 | NA | 49 | NA | 8 | NA | NA | 1 | 1 |
| 47 | Control | 2 | NA | 44 | NA | 4.1 | NA | NA | 1 | 1 |

### Animal Model

A mouse line was generated carrying the E280A mutation, C57BL/6-Psen1em1(E280A) (hereafter referred to as PSEN1Ki mouse, Suppl. Fig. 1). The mouse was generated by CRISPR/Cas in a C57/BL/6 background and did not express any deleterious phenotype for at least 24 months of age. Animals were aged to the experimental time points and brains were collected for formalin fixation and paraffin embedding, snap-frozen storage, or transcardiac perfusion with paraformaldehyde/glutaraldehyde for electron microscopy (EM) by transcardiac perfusion with paraformaldehyde/glutaraldehyde (see detailed methods below).

### Histological and immunohistochemical analysis

In this study brain tissues from Brodmann areas 10 (frontal cortex – FC) and 17 (occipital – OC) were evaluated with the intent that the greater CAA laden OC could also be compared with the FC, a region which usually has lower burden of CAA. Four µm thick sections were, as previously published^9^, stained for haematoxylin and eosin (H&E) and further processed for immunohistochemical (IHC) staining for amyloid beta (Aβ, 1:100, Mob410; Diagnostic BioSystems Inc., Pleasanton CA, USA), Tau (AT8, 1:200, MN1020, Thermo Fisher Scientific Inc., Waltham MA, USA,), glial fibrillary acidic protein (GFAP, 1:200, M0761, DAKO GmbH, Jena, Germany), Ionized calcium-binding adapter molecule 1 (Iba1, 1:500, 019-19741; FUJIFILM Wako Chemicals Europe GmbH, Neuss, Germany), Transmembrane Protein 119 (TMEM119, 1:100, <u>400102</u>; Synaptic Systems GmbH, Göttingen, Germany). Automatic immunostaining was performed with a Ventana Benchmark XT system (Roche, Basel, Switzerland) according to manufacturer instructions. Briefly, after dewaxing and inactivation of endogenous peroxidases with 3% H2O2/PBS, antibody specific antigen retrieval was performed, sections were blocked and afterward incubated with the primary antibody. For detection of specific binding, the ultraView Universal DAB Detection Kit (Ventana Medical Systems, Inc., Tucson, Arizona, USA) containing secondary antibodies, DAB stain and counter staining reagent was used. We also performed Elastica van Giesson staining in FC and OC from both AD variants. Stained sections were scanned using a Hamamatsu NanoZoomer S20 Digital slide scanner (Hamamatsu Photonics, Hamamatsu, Japan) and images of whole stained sections were obtained at a resolution of at least 1 pixel per µm. Total signal measurements of Aβ, Tau, Iba1 and TMEM119 were performed with ImageJ Fiji Software (version 1.52p, NIH, Bethesda, MA, USA)^11^.

### Assessment of vascular pathology

Vascular pathology was assessed as previously described using the Deramecourt V et al. and the vascular cognitive impairment neuropathology group (VCING) scales^9^. Briefly, evaluated features included arteriosclerosis, CAA, perivascular hemosiderin leakage, PVS dilatation and myelin loss. The presence of cortical microinfarcts, large or lacunar infarcts and haemorrhages were considered for the final scoring.

### Quantification of CAA

CAA was reliably quantified as described before^9^. Leptomeningeal vessels, cortical vessels and capillaries were graded for Aβ immunoreactivity and scores were given based on severity. Capillary CAA was evaluated as present or absent. Additionally, the total number of cortical and capillary CAA affected vessels was divided by the total number of unaffected vessels.

### PVS quantification

PVS quantification was performed as previously described^8^. In brief, the ImageJ Fiji Software was used to measure the longest distance between the parenchyma and the vessel to determine the exact size of PVS. The PVS ratio was determined by dividing the measured distance by the diameter of the measured vessel. The PVS dilatation was quantified in the OC of 32 FAD and 10 SAD cases, and in the FC for 20 FAD cases and 10 SAD cases.

### Serial Block-face Scanning Electron Microscopy (SBEM)

Brain slices were prepared for SBEM as previously described^12,13^. Briefly, mice were anesthetized with an intraperitoneal injection of ketamine/xylazine and were transcardially perfused with a brief flush of Ringer’s solution containing heparin and xylocaine, followed by approximately 50 mL of 2.5% glutaraldehyde/2% paraformaldehyde in 0.15 M sodium cacodylate buffer containing 2 mM CaCl2 (CB, pH 7.4). The brain was post-fixed overnight on ice in the same fixative and cut into 100 µm thick coronal sections. Then, brain slices were washed with 0.15 M CB and placed into 2% OsO4/1.5% potassium ferrocyanide in 0.15 M CB containing 2 mM CaCl2 for 1 h at room temperature (RT). After thorough washing in double distilled water (ddH2O), slices were placed into 0.05% thiocarbohydrazide for 30 min. Brain slices were again washed and then stained with 2% aqueous OsO4 for 30 min. Subsequently, brain slices were washed and placed into 2% aqueous uranyl acetate overnight at 4°C. Following this, brain slices were washed with ddH2O at RT and then stained with 0.05% en bloc lead aspartate for 30 min at 60°C. After another round of washing with ddH2O, brain slices were dehydrated on ice in 50%, 70%, 90%, 100%, and 100% ethanol solutions for 10 min at each step. Brain slices were then washed twice with dry acetone and placed into a 50:50 mixture of Durcupan ACM and acetone overnight. Subsequently, brain slices were transferred to 100% Durcupan resin overnight and then flat-embedded between glass slides coated with mold-release compound, left in an oven at 60°C for 72 h. SBEMs were performed using Gemini SEM (Zeiss, Oberkochen, Germany) equipped with a Gatan 3 View system and a focal nitrogen gas injection setup. This system allowed precise application of nitrogen gas over the block face of ROI during imaging with high vacuum to maximize SEM image resolution and ensure reliable sectioning. Images were acquired with a 2.5 kV accelerating voltage and a 1 µs dwell time, with XY pixels of 4 or 5 nm, 60 nm Z steps, and a raster size of 15 k × 15 k in x and y. Volumes were collected using a 65% to 85% nitrogen gas injection under high vacuum. After the collection of volumes, the histograms for the slices throughout the volume stack were normalized to correct for drift in image intensity during acquisition. Volumes were aligned using cross correlation, segmented, analysed and visualized using IMOD^14^.

### Vessel isolation for proteomic analysis and immunofluorescence

Approximately 400 mg of cortical tissue from 20 frontal cortices (FAD = 10, SAD = 5, Healthy Controls = 5) and 30 occipital cortices (FAD = 10, SAD = 5, Healthy Controls = 5), were homogenized in MCDB131 medium (#10372019, Thermo Fisher Scientific Inc., Waltham MA, USA) containing protease and phosphatase inhibitor (#04693159001 and #04906837001 respectively; Roche, Basel, Switzerland) and centrifuged (4°C) at 2000 xg for 5 min. The resulting pellet was resuspended in 15 % (wt/vol) 70-kDa dextran and centrifuged (4°C) for 15 min at 10.000 g.

In the case of peptidomics, the microvessel-containing pellet was retrieved and resolved in 300 µl lysis buffer (150mM NaCl, 10mM NaH2PO4, 1% Triton X-100, 0.5% SDS, 0.5% Deoxycholate), sonicated and centrifuged (4°C) at 14000 xg for 15 min. 40 µl of the vessel lysate was transferred to a new tube for total vessel proteomics. The rest of the supernatant was used for immunoprecipitation of Aβ using DynabeadsTM Protein G Immunoprecipitation Kit (10007D, Thermo Fisher Scientific Inc., Waltham MA, USA), alongside two antibodies recognising the Aβ 1-16 (6E10, #803003, BioLegend, San Diego CA, USA) antibody and the Aβ 17-24, (4G8, #800703, BioLegend, San Diego CA, USA). All samples were stored at −80°C until analysis.

For immunofluorescence, microvessel-containing pellets were transferred to a 100 µm cell strainers and washed with PBS. Cerebral microvessels trapped on the filter were fixed with 4% PFA/PBS for 15 min. Subsequently, the microvessels were washed with PBS, retrieved in 1% BSA/PBS and centrifuged at 2000 g for 10 min at 4°C. The pellet was resuspended in PBS and 50µl of microvessel solution was placed onto Superfrost microscope slides and air dried. Slides were stored at −80°C until they were stained. The microvessels were moisturised with PBS, permeabilized with 0.25% Triton X-100/PBS and blocked in MAXblockTM Blocking Medium (#15252, Active Motif, Waterloo, Belgium) for 1 hour at RT. Primary antibodies were incubated in MaxBindTM Staining Medium (#15253, Active Motif, Waterloo, Belgium) overnight at 4°C and after washing with MaxWashTM Washing Solution (#15254, Active Motif, Waterloo, Belgium) three times for 5 min each, secondary antibodies were incubated in MaxBindTM Staining Medium for 1 hour at RT. Slides were then washed before mounting with DAPI-Fluoromount-G□(0100-20, SouthernBiotech, Birmingham AL, USA). High resolution images were taken with a Leica TCS SP8 confocal laser scanning microscope (Leica Microsystems, Mannheim, Germany) using a 63X immersion oil lens objective.

### Analysis of the proteome

#### Aβ peptidomics

The raw data were analysed using Proteome Discoverer 2.4. Label-free quantitation (LFQ) was enabled in the processing and consensus steps, and spectra were matched against the Uniprot Homosapiens database, including reviewed and unreviewed proteins. The enzyme was set to “No enzyme (unspecific)”. Dynamic modifications were set as Oxidation (M) and Acetyl on protein N-termini. All results were filtered to a 1% FDR, and protein quantitation done using the built-in Minora Feature Detector.

Next, the peptide readout was analysed using protti^15^, an R package for data analysis of peptide bottom-up proteomics data. Briefly, the peptides intensities dataset was filtered for only those related to Amyloid-beta precursor protein (APP). First, the different types of missingness present in the dataset were defined (complete, missing at random (MAR) or missing not at random (MNAR)). Next, differential analysis on peptide level was performed using a moderated t-test based on the limma R package.

#### Total vessels proteomics

The total vessel proteomics from human samples were analysed using MaxQuant^16^. TMT reporter ion quantitation was enabled in the processing and consensus steps, and spectra were matched against the Uniprot *Homo sapiens* fasta protein database (dated 04-2023). Dynamic modifications were set as Oxidation (M) and Acetyl on protein N-termini. Cysteine carbamidomethyl (on C residues) and TMT 16-plex (on peptide N-termini and K residues) were set as static modifications. All results were filtered to a 1% FDR.

Subsequently, the proteins were also quantified and normalised based on PSMs by using MSstatsTMT^13^. The analysis was performed using the MSstatsTMT 2.8.0 in R version 4.2 with Bioconductor 3.14. First, PSMs were extracted and embedded with the annotation file including TMT related parameters such as run, channel, technical replicate mixture, channel, condition, mixture, and bioreplicates. The embedded data was used as input data and applied proteinSummarization function for MSstats spectrum-level normalization, protein summarization, and protein-level normalisation. After quality control steps, two samples were identified as outliers. For total vessel proteomics from murine samples, microvessels were isolated from whole brains, 9 brains (Controls = 5 and PSEN1Ki = 4) were analyzed. Samples were prepared using the Single-pot, solid-phase-enhanced protocol^17^, and LC-MS/MS measurements were performed using a Orbitrap Exploris 480 MS system (Thermo Fisher Scientific, Waltham MA, USA). Protein identification was conducted using the mouse database and the Sequest algorithm with the Proteome Discoverer Software (Thermo Fisher Scientific, Waltham MA, USA). Data was log2-transformed and normalized across columns. From human and murine samples, proteins with less than 50% missing values condition-wise were used, and the remaining missing values were imputed using median values. The normalized protein summaries were after used as input to empirical Bayes moderated t-statistic using limma^18^. Differential expression is presented as volcano plots that were generated with the ggplot2 package in R.

### Weighted Correlation Network Analysis

Total proteomics data were split based on the region of the cortex and the WGCNA algorithm was used to find correlation with AD types. The WGCNA::blockwiseModules() parameters were as follows: minimum module size of 30, deepSplit = 4, soft threshold power β = 10, merge cut height of 0.25 and signed network. Module eigenproteins were defined, which represent the most representative abundance value for a module, and explain covariance of all proteins within the module. Pearson correlations between each protein and each module eigenprotein were performed; this module membership measure is defined as kME. After the initial network construction, 21 modules involving both AD groups and brain areas, and18 modules for FAD analysis, consisting of 30 or more proteins were detected.

### Gene Ontology Enrichment Analysis

The differential expression proteins (DEP) from each comparison were used for Gene Ontology functional enrichment using clusterProfiler. To clarify the mechanisms underlying the impact of module genes on correlative clinical features, genes in modules of interest were used for Gene Ontology enrichment analysis using the clusterProfiler package^19^.

#### Cell type deconvolution analysis

Whole proteomic expression data were used to estimate the relative proportions of specific brain vessels cell types using the BRETIGEA R package. The analysis focused on astrocytes, endothelial cells, pericytes and smooth muscle cells. Cell type-specific marker genes were retrieved from the Human Protein Atlas.

### Single nuclei RNAseq data analysis

The single nuclei RNA-seq (snRNA-seq) data were generated as described in the original publication^20^ and retrieved using the GEO accession number GSE222494. The available count matrix was then imported into Seurat package^21^. Data processing and annotation of the major cell types were collected from the original publication, and the astrocytes were extracted for further analysis. For each diagnosis group (HC, SAD, FAD), astrocytes were assigned to subset and re-clustered to check for common and unique signatures between the groups and all cells together. This was further checked with Sankey plots generated using the networkD3 package^22^.

Dotplot heatmaps were generated using astrocytic markers genes for control, SAD and FAD astrocytes. Functional attributions for each cluster according to functional enrichment analysis and gene expression profiles were assigned for each cluster and group.

## Results

### Brain Vascular Pathology

Vascular pathology was assessed in a cohort comprising 32 individuals with FAD and 10 with SAD. Vascular pathologies including arteriosclerosis, PVS, microinfarcts, myelin loss, and CAA (Fig. 1A) were evaluated^23^ and summarized into a single score. Compared to SAD cases there was a significantly higher degree of vascular pathology in the Frontal Cortex (FC) of FAD subjects. On the other hand, the Occipital Cortex (OC) was equally affected by vascular pathology in both FAD and SAD cases (Fig. 1B). Arteriosclerosis was observed across regions without significant differences between FAD and SAD (Fig. 1C). Similarly, no significant differences were found in perivascular scoring between the FAD and SAD cases (Fig. 1D). These findings highlight the presence of vascular pathology inAD, which may go undetected before death by neuroimaging methods. To investigate whether the observed difference in vascular pathology between the FC and the OC in FAD is linked to amyloid pathology, cerebral amyloid angiopathy (CAA) was specifically examined (Fig. 1E). The CAA VCING scale was used to evaluate the presence of leptomeningeal, cortical, and capillary CAA^24^. Similar CAA scores were detected in both cortices, FC and OC, in both AD variants, with a tendency for higher CAA scores in FAD cases in the FC (Fig. 1F). Using Elastica van Giesson staining, we also assessed the build-up of Collagen IV (COL4) fibres in small vessels in FC and OC of AD cases. We found significantly higher COL4 reactivity in FAD cases in both cortices (Fig. 1G). To gain further insight into the CAA pathology in these cases, vessels unaffected by CAA, partially affected vessels, and vessels fully covered by CAA were counted in the FC and OC from FAD cases. A higher percentage of confluent CAA vessels were detected in both cortices, but the differences did not reach statistical difference (Fig. 1H), indicating similar CAA pathology between FAD and SAD from a morphological standpoint.

**Figure 1.**
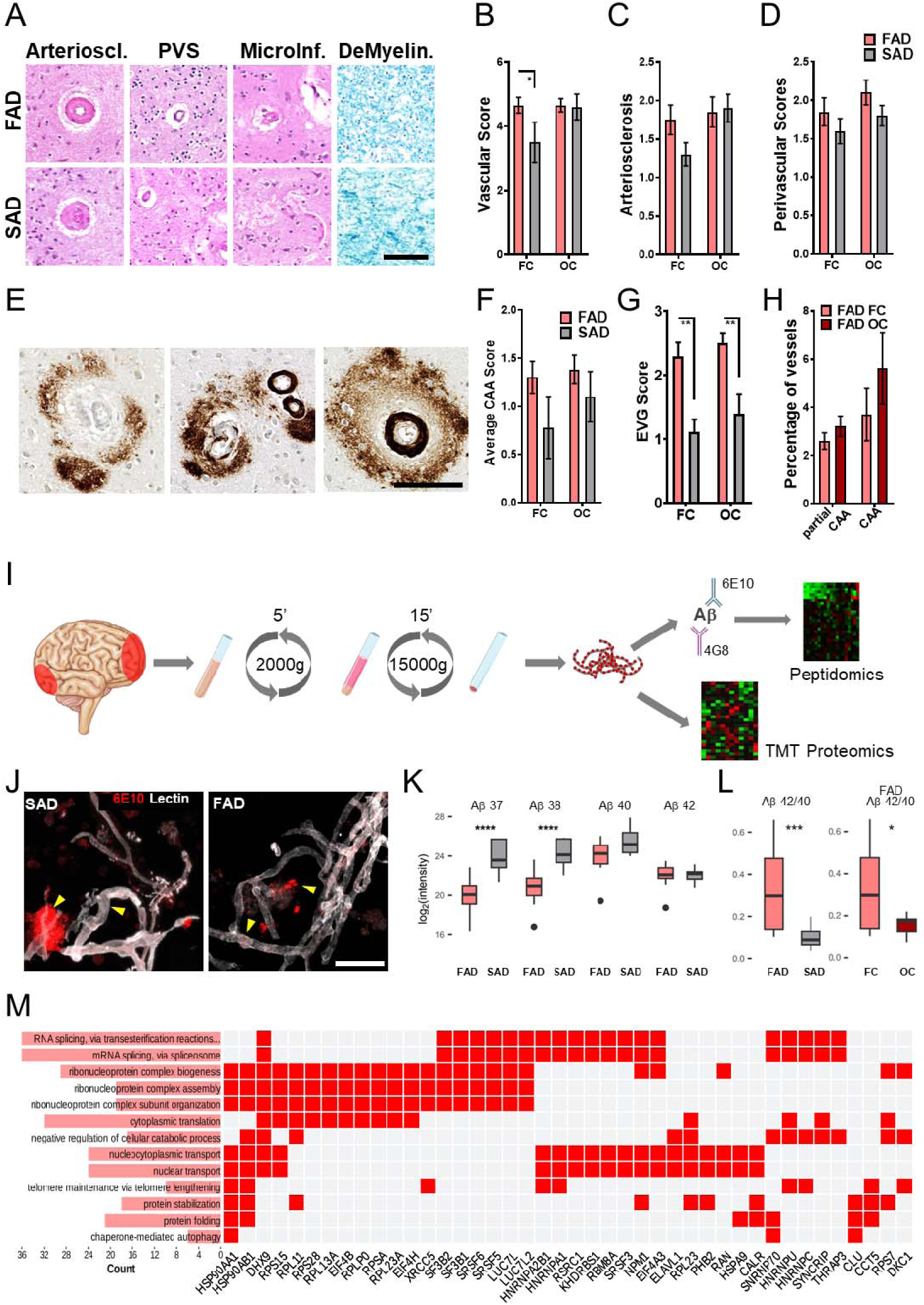
Brain vascular pathology and CAA pathology features in SAD and FAD cases. A. Representative pictures of small vessels disease features in both SAD and FAD cases. Bar = 100 μm. B. Bar graphs for vascular scores in the Frontal Cortex (FC) and Occipital Cortex (OC) of FAD (n = 32) and SAD (n = 10) cases. FC in FAD cases showed to be significantly more affected. C. Bar graphs for arteriosclerosis scores in FC and OC of AD cases, showing no significant differences. D. Bar graphs for perivascular scores in FC and OC of AD cases showing no significant differences. E. Representative pictures for FAD microvessels depicting perivascular diffuse amyloid plaques in absence or presence of CAA pathology. Bar = 100 μm. F. Bar graphs of CAA scores in FC and OC of AD cases, SAD shows a tendency for less CAA pathology in the FC. G Bar graphs for the semiquantitative assessment of Collagen fibres build up in FC and OC of AD cases. FAD vessels showed significantly more Collagen build up in both evaluated cortices. H. Bar graphs for the percentage of vessels partially or totally affected by CAA in the FC and OC of FAD cases. The OC shows a non-significant higher percentage of total CAA vessel pathology. I. Schematics of our experimental design. Briefly, isolated microvessels from FC and OC from AD cases. Protein homogenates were either co-immunoprecipitated with two different antibodies against Aβ for proteomic and peptidomic analyses or directly processed for TMT proteomics. J. Representative images of isolated microvessels from the FC of an SAD and an FAD case., stained for Lectin-488 (white), and Aβ (6E10 antibody, red). Intramural and perivascular Aβ aggregates (yellow arrowheads) can be visualized in both AD variants. Bar = 50 μm. K. Boxplots depicting Ab peptides 1-37, 1-38, 1-40, and 1-42, detected in coimmunoprecipitated microvessels proteins from the FC of FAD and SAD cases. Both shorter Aβ peptide species, 1-37 and 1-38, were significantly lower in FAD. L. Boxplots depicting Aβ 1-42/1-40 ratios in the FC of FAD and SAD, and Ab 1-42/1-40 ratios of FC and OC in FAD cases. FAD FC Aβ 1-42/1-40 ratio was significantly higher than either SAD FC or FAD OC. M. Heat plot of the gene set enrichment analysis (GSEA) of the proteins that coprecipitated with Aβ aggregates in isolated microvessels. Biological processes GO terms, the number of genes detected in each (length of pink bars) and the genes identified are shown. Notably, RNA related functions and protein catabolism terms were identified. p * = >0.05, ** = >0.01, *** = >0.001, **** = >0.0001. Error bars = standard error of mean (SEM). Abbreviations: Arterioscl, arteriolosclerosis; PVS, perivascular space; MicroInf, microinfarct; DeMyelin, demyelination.

### Proteomic analysis of CAA pathology

To understand the potential causes of Aβ-dependent and Aβ-independent vascular pathology and associated structural changes in the vessels of FAD, the proteomes of purified vessels from the FC and OC of AD cases were analyzed (1I). As previously demonstrated in SAD^25^, the presence of Aβ deposits in isolated vessels in FAD and SAD cases was confirmed (Fig. 1J). We then immunoprecipitated, fractions of microvessel protein lysates with two different Aβ antibodies (6E10 and 4G8), and analysed their peptidomic signature to reveal Aβ peptide profiles and associated proteins (Fig. 1I). For this purpose, all Aβ peptides detected were grouped according to their C-terminal ending. For example, Aβ37 contains all peptides ending at position 37, regardless of their total length. Additionally, the Aβ42/Aβ40 ratio was calculated based on the intensities for each peptide. As previously described for Aβ peptides pathological profile in the brains of this FAD population^7^, shorter Aβ peptides species (1-37, 1-38) were more abundant in SAD microvessels from FC (Fig. 1K). As expected, the Aβ42/Aβ40 ratio was higher in FAD than in SAD cases, and for FAD brains, in FC than in OC (Fig. 1L). All the Aβ peptides comparisons are reported in Supplementary data (Suppl. Fig. 2). Moreover, Aβ peptides aggregate into deposits or plaques and CAA. Specific plaque-associated proteins have been previously identified^26^, and it is plausible that similarly co-deposited proteins could be found in Aβ aggregates in microvessels. We analyzed peptides that co-immunoprecipitated with Aβ and performed a functional enrichment analysis, identifying several proteins linked to RNA metabolism, telomere maintainance and protein catabolism, among other biological processes (Fig. 1M).

### Proteomic analysis of isolated brain microvessels

TMT proteomics performed in the remaining fraction of isolated cerebral microvessels from FAD (FC = 10, OC = 20), SAD (FC = 4, O = 5) and healthy control (HC; FC = 5, OC = 5) cases (Fig. 1H) enabled the identification of a total of 5615 proteins. To distinguish true biological differences from Aβ co-aggregating proteins, we removed from further analyses any proteins identified as co-precipitating with Aβ aggregates (Fig. 1L). Unlike previous microvessel proteomic studies, this subtraction ensured that the dysregulated proteins we report reflect genuine biological change rather than passive co-aggregation with amyloid; notably, RNA-processing dysregulation persisted after this removal. Differentially expressed proteins (DEPs) analyses were then conducted in purified vessels from the FC and OC of FAD or SAD versus HC, revealing numerous dysregulated proteins. DEP analyses from FAD, SAD and HC in FC showed similar number of DEP between all three possible comparisons, with the highest number (43 proteins) of common DEP between FAD and SAD in relation to HC, and marginally more unique DEP (144 proteins) identified when comparing FAD and HC (Fig. 2A). On the other hand, when the same comparisons were conducted in OC, FAD showed a higher number of unique DEP when compared to SAD (207 proteins) and HC (496 proteins). Similarly, the highest number of common DEP (145 proteins) was found between FAD and SAD in relation to HC (Fig. 2B). Commonly dysregulated proteins between both AD variants were explored in FC and OC. Several proteins have been previously described in similar proteomic analyses from cerebral microvessels in mild cognitive impairment (MCI), AD and CAA^27,28^, such as SMOC1, APOE and HTRA1 (Fig. 2C-D). Other proteins, such as CLU and RPL23A, co-precipitated with Aβ aggregates. Several of the dysregulated proteins identified in FAD microvessels were glycoproteins. We also performed glycoprotein-targeted proteomic analyses to FC and OC microvessels from HC and FAD cases. Among dysregulated proteins in both cortices, there were 8 proteoglycans, five of them in the OC (Suppl. Fig. 3A). These proteoglycans and some of their interacting partners were mostly upregulated in both cortices (Suppl. Fig. 3B-C). Also, heparan sulfate-binding could be identified among dysregulated associated protein functions in the OC (Suppl. Fig. 3D). These proteins included APOE, SLIT2, RSPO2, ADAMTS1, MDK, VTN, FGF1, APOB and PRELP (Suppl. Fig. 3E).

**Figure 2.**
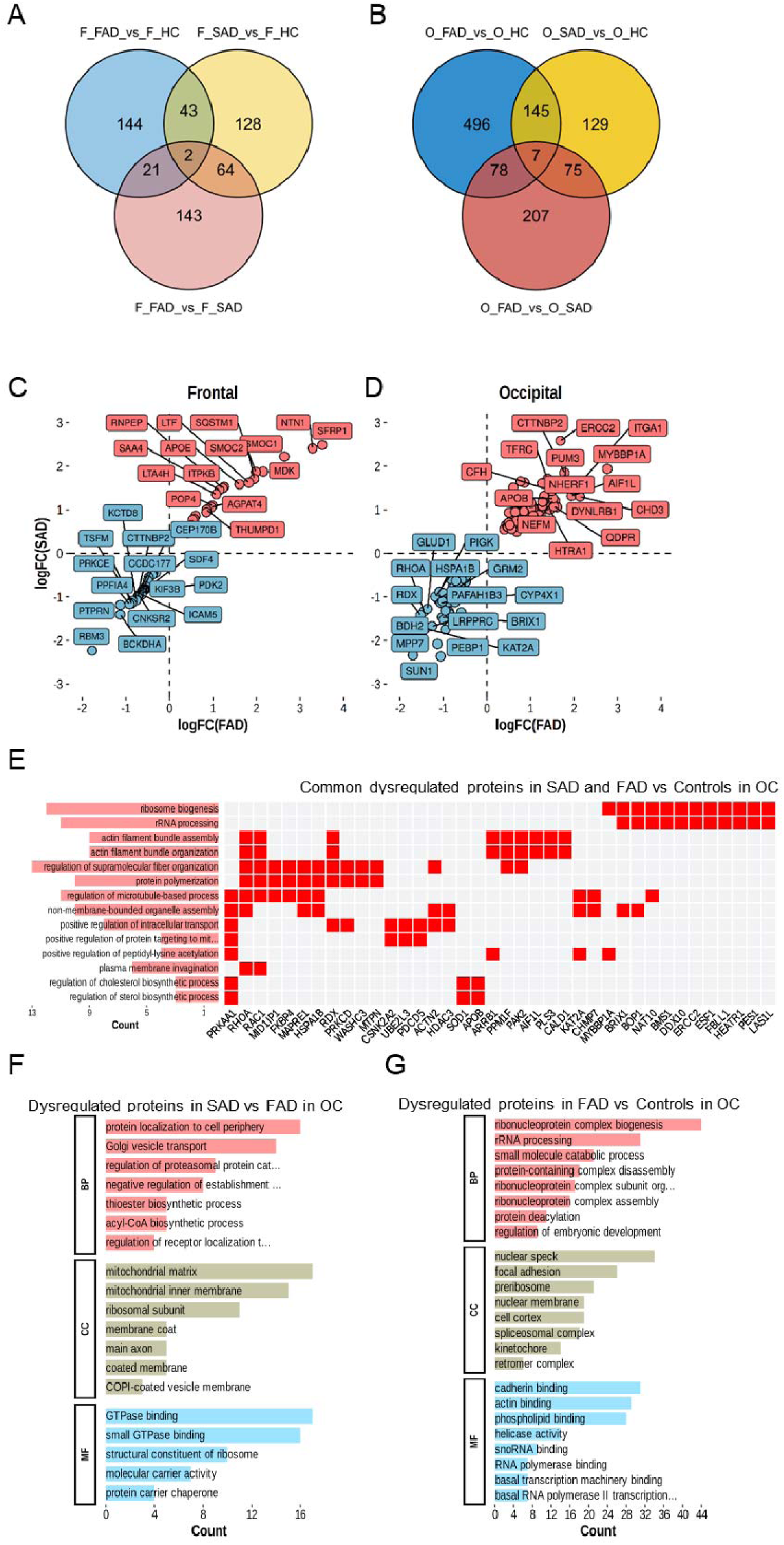
Proteomic analysis of isolated microvessels in FAD and SAD cases. A. Venn diagram depicting DEPs identified in the comparisons between FAD and healthy controls (HC), SAD and HC, and FAD and SAD from FC proteomes. A similar amount of uniquely identified DEPs were detected in the comparison between FAD and each of the two other groups. B. Venn diagram depicting DEPs identified in the comparisons between FAD and healthy controls (HC), SAD and HC, and FAD and SAD from OC proteomes. A remarkable higher number of DEPs were detected in the comparison between FAD and controls. C. Scatter plot for fold change (FC) values of commonly upregulated (red) and downregulated (blue) proteins in SAD and FAD, in the FC. D. Scatter plot for fold change (FC) values of commonly upregulated (red) and downregulated (blue) proteins in SAD and FAD, in the OC. E. Heat plot of the GSEA of the commonly dysregulated proteins in OC isolated cerebral microvessels from bot AD variants. Biological processes GO terms, the number of genes detected in each (length of pink bars) and the genes identified are depicted. Biosynthesis, structural proteins and RNA-associated processes were identified. F. GSEA barplots for biological processes (PB), cellular compartment (CC), and molecular function (MF), for uniquely dysregulated proteins in the comparison between FAD and SAD in the OC. Protein catabolism, mitochondrial membranes, and GTPase binding functions were identified. Notably, some terms included RNA-related proteins. G. GSEA barplots for BP, CC, and MF for uniquely dysregulated proteins in the comparison between FAD and controls in the OC. Mainly RNA-associated proteins were identified among all three term categories.

**Figure 3.**
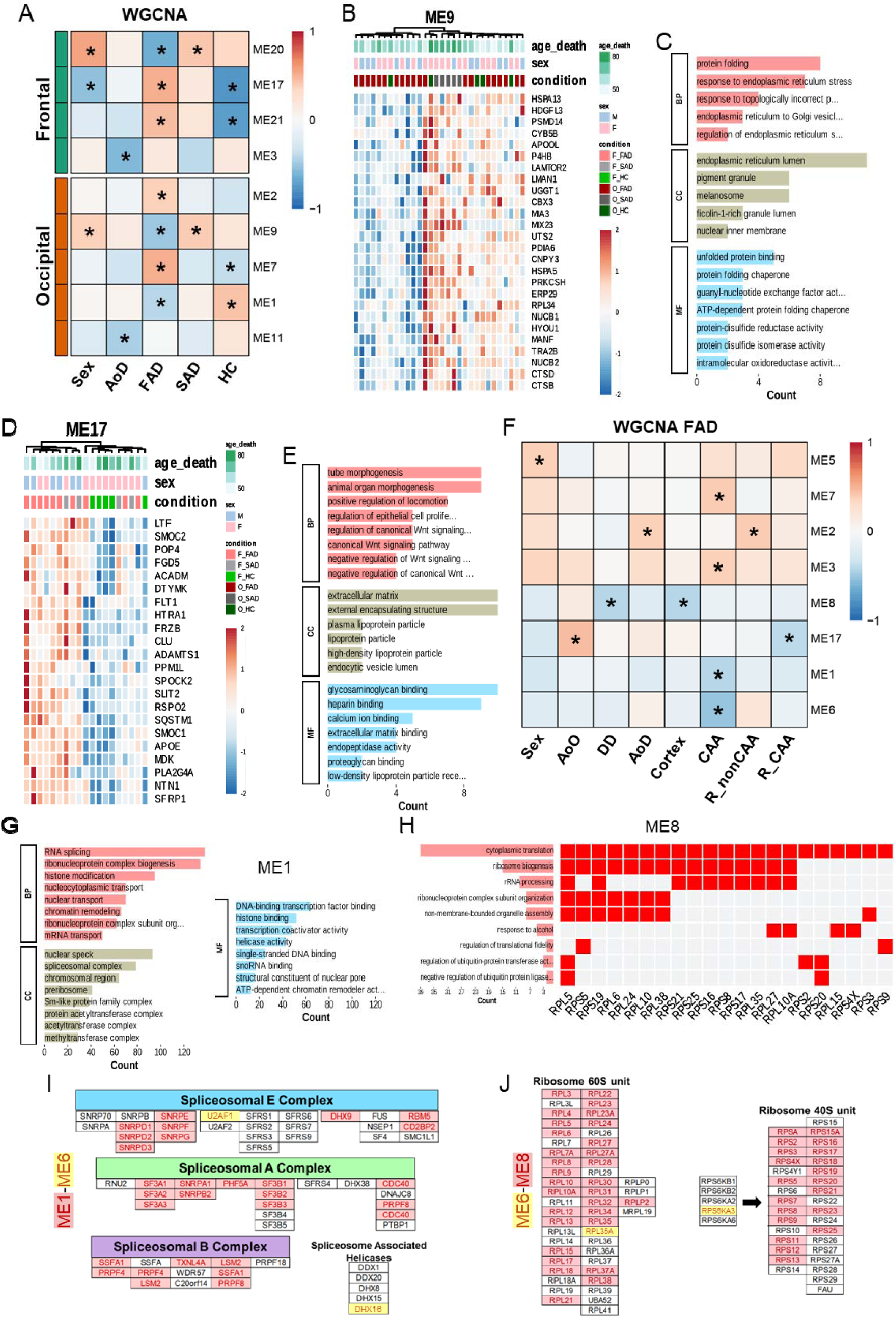
Co-expression networks and associated phenotypes in AD. A. Heatmap of Weighted Co-expression Network Analysis (WGCNA) conducted in the proteome of FC and OC of FAD, SAD cases, and healthy controls (HC), according also to their sex and age of death (AoD). Modules (ME) with statistically significant associations were further labelled with an asterisk. B. Heatmap depicting the proteins belonging to ME9 for the OC, clustered by age of death, sex and condition (SAD or FAD). C. GSEA barplots for BP, CC, and MF for the proteins identified in ME9. Endoplasmic reticulum stress associated processes were identified in this module. D. Heatmap depicting the proteins belonging to ME17 for the OC, clustered by age of death, sex and condition (SAD or FAD). E. GSEA barplots for BP, CC, and MF for the proteins identified in ME17. Structural ECM, WNT signaling and lipid metabolism processes were identified in this module. F. Heatmap of WGCNA conducted in the proteome of FC and OC of FAD, according also to their sex, age of onset (AoO), disease duration (DD), AoD, the severity of CAA pathology (CAA), PVS dilation ratio in vessels without CAA pathology (R_nonCAA), and PVS dilation ratio in vessels with CAA pathology (R_CAA). Modules (ME) with statistically significant associations were further labelled with an asterisk. G. GSEA barplots for BP, CC, and MF for the proteins identified in ME1. Numerous RNA-associated processes were identified in this module. H. Heat plot of the GSEA of the proteins identified in ME8. Biological processes GO terms, the number of genes detected in each (length of pink bars) and the genes identified are depicted. Several RNA-associated processes were also identified in this module. I. Schematic representation of the spliceosomal complex, in its three subunits. Proteins identified in ME1 are labeled in red, proteins identified in ME6 are labeled in yellow. J. Schematic representation of the ribosomal complex, in its two subunits. Proteins identified in ME8 are labeled in red, proteins identified in ME6 are labeled in yellow.

Gene Ontology (GO) enrichment analysis was performed subsequently to determine the biological functions of the dysregulated proteins. Protein polymerization, actin assembly and RNA processing were commonly dysregulated in AD groups versus HC in the OC (Fig. 2E). Additionally, commonly dysregulated proteins in both AD groups in the FC included mainly adhesion and synaptic proteins (Suppl. Fig. 4A-B). In the FC of FAD, cellular components such as adhesion and membranal components, together with oxidoreductase activity and dopamine receptor binding molecular functions were significantly dysregulated in unique DEP (Suppl. Fig. 4C). Similarly, significant cellular components such as recycling endosomes and glycoproteins were identified in unique dysregulated proteins in the FC of SAD (Suppl. Fig. 4D). In general, the FC of FAD presented with a more unique proteomic signature. When FAD and SAD OC proteomes were compared, protein biosynthesis was found to be dysregulated, involving mitochondrial and cell membrane proteins (Fig. 2F). Finally, even though proteins associated to RNA related processes and structures were commonly dysregulated in the OC of both AD variants, additional proteins related with these functions were identified dysregulated only in FAD cases in this brain region (Fig. 2G). These findings suggest distinct pathological signatures between the two regions, with the FC exhibiting unique proteomic alterations, and strong dysregulation of RNA associated proteins in FAD.

Next, a weighted gene co-expression network analysis (WGCNA) was performed to study the relationship between proteins present within all samples. Across all three network analyses, two functional themes recurred and tracked disease: modules enriched for RNA and DNA processing, which correlated negatively with FAD and CAA severity, and modules enriched for extracellular matrix and collagen organization, which positively correlated with FAD and PVS dilation. Three separate WGCNA analyses were conducted: one for all cases from the FC, one for all cases from the OC (Fig. 3A-E); and one limited to FAD cases (Fig. 3F-J). For the first analysis, the significant modules identified were correlated with sex, age of death (AoD), and the groups (FAD, SAD, and HC). The largest module, ME1 was enriched in biological processes related to RNA processing and DNA repair, with molecular functions associated with DNA and mRNA binding. ME2, positively correlated with FAD, was associated with synaptic organization, actin filament organization, and neuron projection morphogenesis, with cellular components related to the cell cortex and actin cytoskeleton. Modules related to cellular respiration, oxidative phosphorylation, and mitochondrial function were negatively correlated with AoD (ME3 and ME11). Conversely, ME7 positively correlated with FAD, was enriched in processes related to morphogenesis and cell migration, with components related to the ECM and binding functions. In the OC, ME9 exhibited a significant negative correlation with FAD, as well as negative correlations with sex and SAD. This module was enriched in biological processes related to the regulation of response to endoplasmic reticulum (ER) stress and protein folding, with cellular components related to pigment granules and melanosomes. Molecular functions identified included protein disulfide isomerase activity and intramolecular oxidoreductase activity (Fig. 3B-C). ME11, negatively correlated with AoO and FAD (Fig. 3A), showed significant enrichment in biological processes related to collagen fiber organization, with cellular components associated with collagen-containing extracellular matrix. Molecular functions enriched in this module included collagen binding, ECM structural constituent, and extracellular matrix binding. ME17 positively correlated with FAD and negatively correlated with healthy cases and sex in the FC. It revealed proteins mainly involved in the regulation of the Wnt pathway and ECM. Notably, it also included key AD related proteins and was already detected in abnormal AD microvessels^27,28^ and mentioned in Fig. 2C is commonly dysregulated in both AD variants, such as SMOC1, SMOC2, APOE, CLU and HTRA1 (Fig. 3D-E). ME20 contained proteins primarily associated with mitochondrial function, such as OPA3 involved in mitochondrion organization and SLC25A1 involved in citrate transport across the inner mitochondrial membrane. Other proteins in ME20 had diverse functions, including regulation of endothelial cell proliferation (EDF1) and GTP binding activity (ARL15). ME21 correlated positively with FAD and negatively with healthy controls. These findings highlight diverse molecular pathways and functions represented within different WGCNA modules, providing further insights into the complex interplay of biological processes underlying AD pathology in different cortical regions.

To further explore the biological disparities between the FC and OC within the FAD group, WGCNA was repeated using all the FAD samples (Fig. 3F). This analysis included proteins specific to FC and OC, allowing for the identification of modules correlated with various clinical and pathological features. ME1 negatively correlated with CAA, and its proteins were associated with RNA and DNA processing (Fig. 3G). The next largest module, ME2, positively correlated with AoO and the ratio of PVS in non-CAA vessels, and it was enriched in processes related to vesicle-mediated transport and protein folding, with cellular components related to vesicle tethering complexes and endocytic vesicles. ME3 positively correlated with CAA, showing an enrichment in terms related to structural organization and ficolin-1-rich granule lumen, while molecular functions included fatty acid binding and actin filament binding. ME5, at its turn, positively correlated with sex, was enriched in processes related to ATP metabolism. ME6 negatively correlated with CAA, was enriched in processes related to RNA and DNA processing. ME7 positively correlated with CAA, exhibited enrichment in immune response pathways and ECM components. Molecular functions enriched in this module included Heparin and ECM binding (Suppl. Fig. 5). ME8 negatively correlated with duration of disease (DD) and FC and it was associated with proteins involved in RNA processing and ribosome assembly (Fig. 3H). To note, modules ME1, ME6 and ME8 included a large number of proteins related to RNA regulation and metabolism (Fig. 3G, H), specifically, proteins involved in spliceosome (Fig. 3I) and ribosome (Fig. 3J) assembly.

Finally, ME17 positively correlated with AoO and negatively correlated with the ratio of PVS in CAA-affected vessels, containing processes related to extracellular remodelling, such as collagen fiber organization, angiogenesis, and blood vessel development. Significant cellular components in this module included collagen-containing ECM and protein complexes involved in cell-matrix adhesion (Fig. 4A). To validate the proteomics findings, different markers were selected for immunofluorescence studies. Secreted modular calcium-binding protein 1 (SMOC1) was selected based on previous proteomic studies^27,28^ and its colocalization with Aβ plaques and CAA in the current study. Immunofluorescence staining revealed more intense SMOC1 immunoreactivity in the arterioles of FAD when compared to SAD. However, there was virtually no reactivity in Aβ deposits or plaques (Suppl. Fig. 6A). Proteomic analysis of purified vessels showed, SMOC1 to be more abundant in FAD, than in SAD but the difference was not significant(Suppl. Fig. 6B). Our findings identified several ECM glycoproteins. Tenascin-C (TNC) was selected for validation due to its reported production by astrocytes, while versican (VCAN) has been shown to be produced by endothelial cells. Similarly, VCAN exhibited comparable signal intensity between AD groups, with no detectable signal in COL4-positive areas (Suppl. Fig. 7A). Levels of TNC in the proteomic analysis did not differ between FAD and SAD, which was consistent with the immunostaining results (Suppl. Fig. 7B). Therefore, quantitative validation of the extracellular matrix signature rested instead on the markers that reached significance — increased collagen build-up, loss of COL4 coverage, and increased vascular tortuosity and lacunarity in FAD — while SMOC1, TNC and VCAN staining served to confirm vascular localization rather than differential abundance.

**Figure 4.**
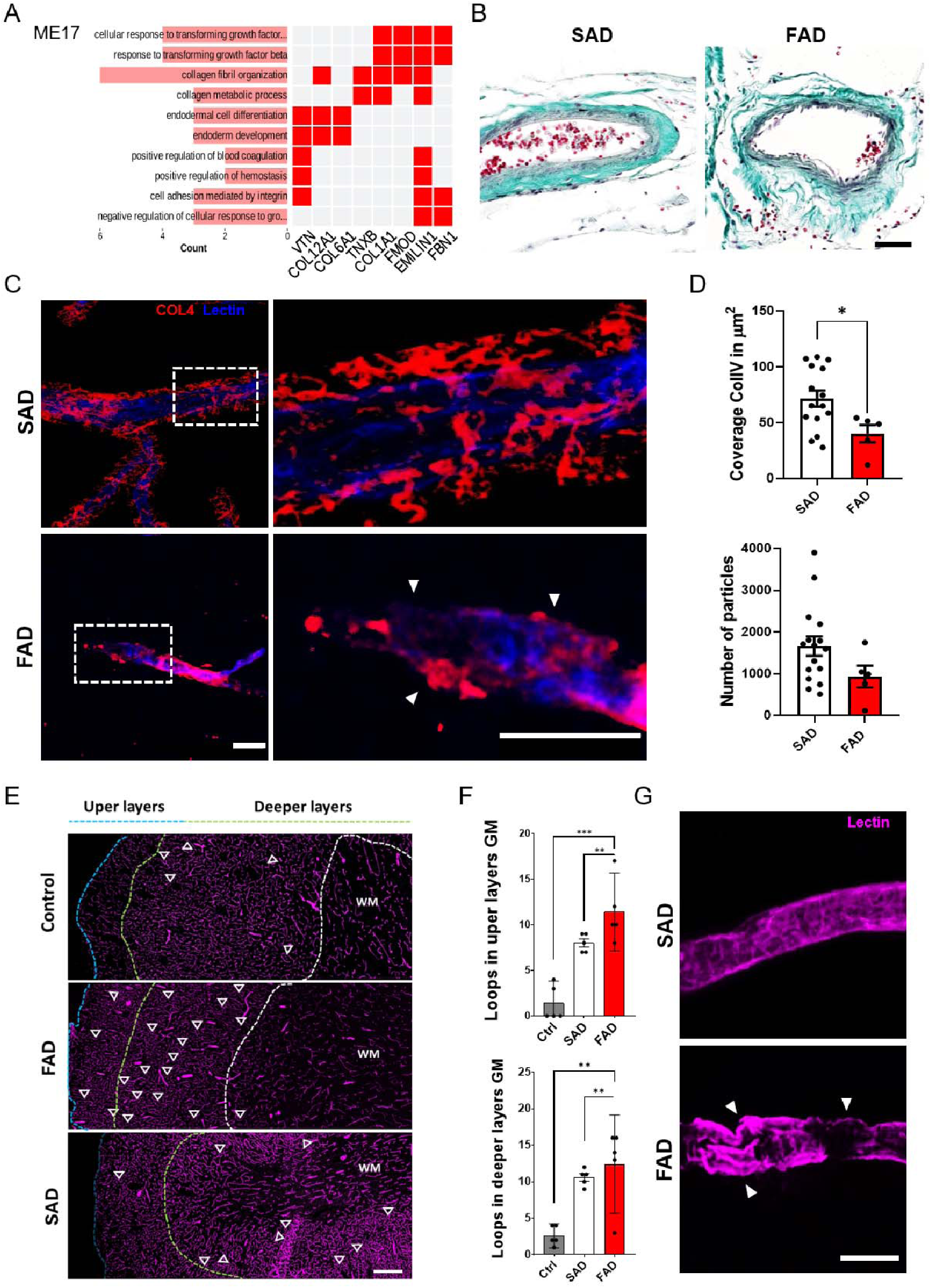
Abnormal Extracellular Matrix (ECM) and vascular networking in PSEN1 FAD cases. A. Heat plot of the GSEA of ME17 proteins from FAD variants. Biological processes GO terms, the number of genes detected in each (length of pink bars) and the genes identified are depicted. Extracellular matrix proteins were identified. B. Representative pictures of Masson’s trichrome staining of leptomeningeal vessels of both AD variants. FAD vessels show disorganized collagen in the adventitia layer. Bar = 50 μm. C. Representative images of Collagen IV (COL4, red) and Lectin-488 (blue) staining of cerebral microvessels from both AD variants. FAD vessels show Collagen loss and uneven coverage (white arrowheads). Bar = 20 μm. D. Barplots for COL4 coverage and number of COL4 positive particles in microvessels from SAD and FAD cases. COL4 vessel coverage was significantly lower in FAD. E. Representative images of Lectin staining of cortical and white matter (WM, white dashed line) microvascular network in upper cortical layers (blue dashed line) and deeper cortical layers (green dashed line). Abnormal loops are depicted as dashed white circles. Bar = 50 μm. F. Bar plots for abnormal vascular network loops identified in upper and deeper layers from the FC of both AD variants. FAD cases showed a significantly higher number of abnormal loops. G. Magnified representative pictures from E of individual microvessels. FAD microvessels are tortuous and show knots (white arrowheads). Bar = 10 μm.

The findings from ME17 align with previous observations of ECM-related changes in the PVS in the FC of FAD cases^9^, highlighting the importance of extracellular remodelling in the pathology of FAD. To validate this observation, Masson-Goldner staining was performed, revealing collagen disorganization in the meningeal large vessels of FAD cases, while meninges in SAD cases appeared intact (Fig. 4B). Furthermore, we immunostained FC microvessels in both SAD and FAD cases and identified significantly increased loss of COL4 coverage in FAD (Fig. 4C-D). Additionally, FAD microvessels showed increased tortuosity and kinking when compared with SAD cases (Fig. 4E-G). These analyses further support the association between ECM dysregulation, abnormal angiogenesis, and blood vessel fragility in FAD cases. Notably, cell deconvolution analysis indicated that astrocytes were the primary contributors to the proteome in FAD cases (Suppl. Fig. 8).

Summarizing our main findings, the WGCNA highlighted different protein profiles associated with AD type, sex and AoO. FAD subjects positively correlated with protein modules involved with structural processes in the OC, and processes and proteins already reported for AD^13,14^ in the FC. A protein module associated with DNA/RNA processes correlated negatively with FAD and differentiated them from healthy controls in the OC. SAD subjects correlated positively with ER stress / protein folding proteins module, while FAD correlated negatively with it. When the WGCNA analysis was focused only on FAD and clinical or vascular pathology features we previously described in this population^9^, we found that CAA pathology is a dominant feature and positively correlated with proteins associated with catabolic and structural functions. Furthermore, another relevant vascular pathology feature in FAD, the increase of PVS, either in CAA unaffected vessels or CAA affected vessels, correlated either with endocytic associated proteins or with ECM proteins, respectively. This involvement of the ECM in vascular pathology in FAD was further validated with the identification of disorganization and abnormal collagen coverage, together with increased vascular tortuosity and vascular network abnormalities. Finally, CAA negatively correlated with RNA or DNA metabolism and processing. In fact, proteins associated with these processes were also negatively correlated with disease duration and cortical area. Throughout the proteomic analysis of AD microvessels, we identified proteins involved in RNA metabolism and function at several instances, either co-precipitating with A peptides, commonly dysregulated in AD cases when compared to controls, or as a consistent proteomic signature of SVD in FAD cases and associated with pathological features.

### Assessment of SVD pathology in an PSEN1 E280A knock-in murine model

SVD presents with multiple features in AD, evident from imaging and neuropathological analysis^8^. Given the multifactorial nature of this disorder, it is perhaps not surprising that when a PSEN1 mutation is the underlying cause, several protein networks appear to be involved and associated with different clinical or pathological features. However, since postmortem analysis of AD brains is usually undertaken in a terminal stage of disease, it was not possible to differentiate which protein networks and molecular functions are a consequence of the pathological aggregation of AD proteins and which are directly modulated by dysfunctional PSEN1. To this end, we generated a PSEN1E280A (PSEN1Ki) transgenic mouse. Because this knock-in carries no human APP and develops neither plaques nor tangles, any pathology it shares with human FAD can be attributed to PSEN1 dysfunction rather than to amyloid or tau aggregation. Homozygous mice do not show a visible deleterious phenotype in their first year of life. However, 3D analysis of the vascular networks in this model showed that cerebral microvessel structure in 6 months old and one year old animals (Fig. 5A) are significantly increased in both, branch length and the branch length-to-number ratio, compared to wild-type animals (Fig. 5B). An increased vascular tortuosity in the cortex of 6 months old PSEN1Ki mice was also found but, not visible at 12 months of age (Suppl. Fig. 9). We isolated cerebral microvessels of PSEN1Ki and wild type mice and performed a similar proteomic analysis as the one conducted in the human samples. We identified differentially expressed proteins involved in processes such as vesicle transport, catabolism, and notably, RNA metabolism and transcriptional regulation (Fig. 5C-D). Given that RNA-related proteins were also detected in PSEN1E280A FC and OC microvessels, we compared the functional gene sets detected in both, PSEN1E280A human microvessels and PSEN1Ki microvessels. In effect, we found RNA-related biological processes commonly dysregulated in both organisms, including ribonucleoprotein complex biogenesis and RNA processing, some exclusive to human FAD microvessels such as ribosome biogenesis, and some exclusive to murine PSEN1Ki microvessels such as protein-RNA complex assembly (Fig. 5E-F). This finding suggests that among the possible multiple factors influencing SVD in PSEN1 E280A cases, at least one factor is directly related to PSEN1 dysfunction due to the E280A mutation.

**Figure 5.**
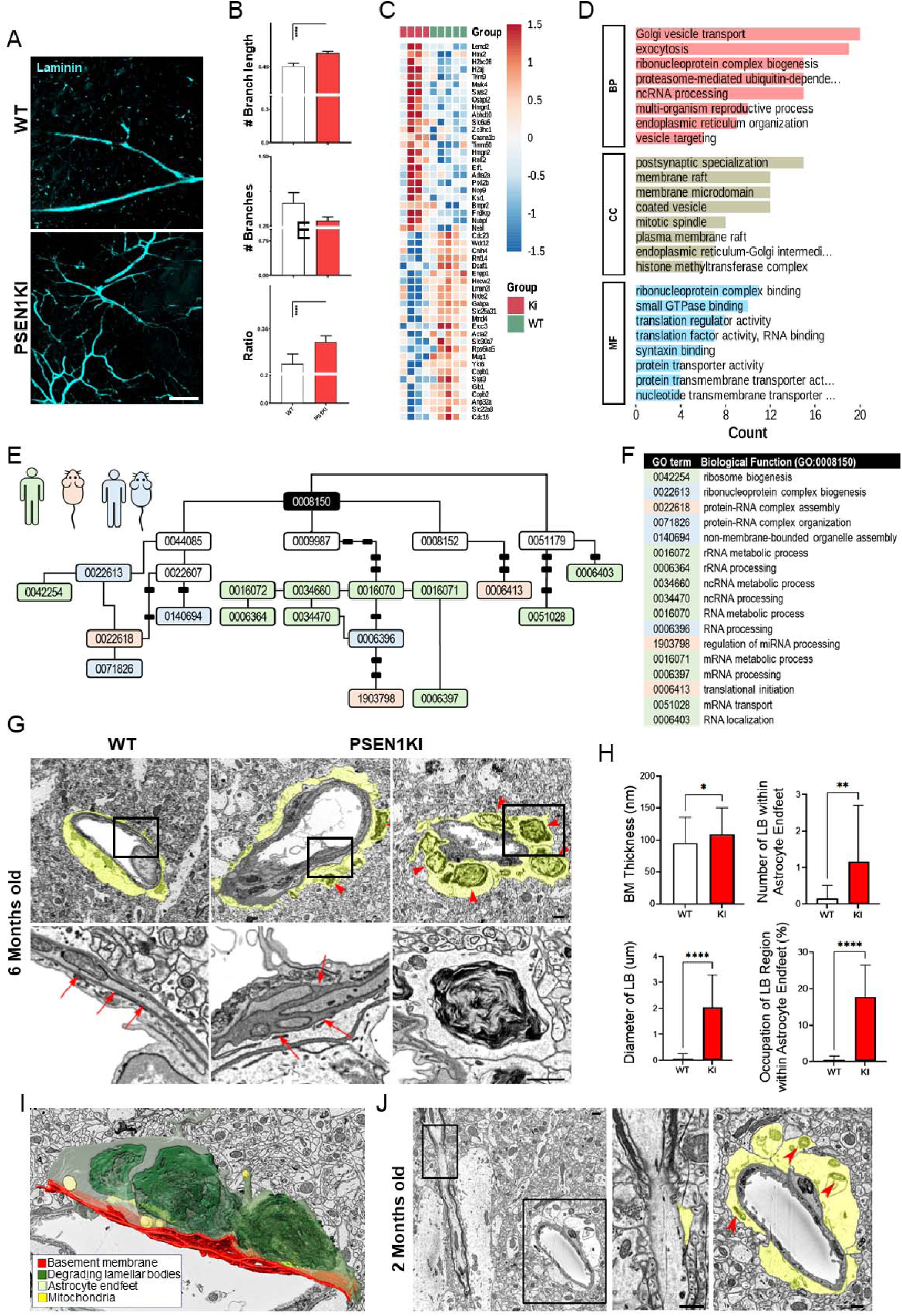
Abnormal microvascular phenotype in PSEN1Ki E280A (PSEN1Ki) mice. A. Representative images in 2D of clarified brain tissue from wild type (WT) and PSEN1Ki 6 months old mice with microvessels stained with Laminin. Bars = 200 μm B. Barplots of three-dimensional analyses of vascular branches length, number, and length/number of branches ratio. PSEN1Ki mice showed significantly longer branches and length/number ratio. C. Heatmap of the DEPs identified in the proteomic analysis of PSEN1Ki (n=4) and WT (n=5) isolated microvessels. D. GSEA barplots for BP, CC, and MF for the DEPs identified in C. Several BP and MF were associated with RNA-related processes. E. Scheme of the GO terms identified as dysregulated in human FAD microvascular proteome (pale green), PSEN1Ki microvascular proteome (pale red), and both (pale blue). F. Table of the GO terms description depicted in E. G. Representative low magnification electron micrographs, and lower magnified panels (black squares) of microvessels from 6 months old WT and PSEN1Ki mice. Astrocytic end-feet are shadowed in pale yellow. PSEN1Ki microvessels show abnormal degrading lamellar bodies in astrocytic end-feet (red arrowheads) and thickening of the basement membrane (red arrows). Bars = 500 nm. H. Barplots for basement membrane (BM) thickness, number of lamellar bodies (LB) within astrocytic end-feet, diameter of LB, and percentage occupation of LB region within astrocytic end-feet. PSEN1Ki mice showed higher values in these variables. I. 3D reconstruction generated from SBEM data of a 6-month-old PSEN1Ki mouse, highlighting BM (red), degrading lamellar bodies (green), astrocytic endfeet (light green), and mitochondria (yellow). J. Representative low magnification electron microphotographs of an axon and a microvessel from a 2-month-old PSEN1Ki mouse, astrocytic processes are shadowed in pale yellow. A small astrocytic process extends onto the nodal membrane, a region where astrocytes typically form fine processes in contact with CNS myelinated nerve nodes. In contrast, the adjacent perivascular astrocytic end-feet appear swollen. Small electron-dense degrading LB (red arrowheads) are evident, suggesting early pathological changes affecting Blood–Brain Barrier (BBB) integrity as early as 2 months. Bars = 500 nm.

To gain understanding of the impact of the PSEN1E280A mutation in brain vessels, we studied the ultrastructure of microvessels in the hippocampi of 6 months old PSEN1Ki mice, compared against wild type controls. We found significant thickening of the basement membrane in PSEN1Ki mice (Fig. 5G-I). In addition, we identified the accumulation of electron-dense material within perivascular astrocytic end-feet, morphologically resembling degrading lamellar bodies, possibly of lysosomal origin^29^. These lamellar bodies were significantly more numerous, larger, and occupying more end-feet in the vicinity of PSEN1Ki microvessels (Fig. 5G-I). These findings suggested a chronic process affecting perivascular astrocytes in PSEN1 mutants. When the microvessel ultrastructure of two months old mice were assessed, similar smaller degrading lamellar bodies material was identified in perivascular astrocytic end-feet, which appeared swollen, suggesting early pathological changes affecting Blood-Brain Barrier integrity as an early phenotypic trait in PSEN1Ki mice (Fig. 5J).

### Transcriptomic evidence of vessel associated astrocytic pathology in FAD

We have previously reported abnormal PVS and lower levels of AQP4 in FAD cases, indicating glymphatic dysfunction and astrocytic involvement in PSEN1 E280A FAD cases^9^. Subsequently, we reported postmortem findings in a resistant homozygous APOE Christchurch FAD case carrying the PSEN1 E280A mutation. One factor that was shown to be associated with the resistance to pathology, and evidenced by morphological and single nuclei transcriptomic findings, was the presence of homeostatic hyporeactive astrocytes^30,31^. We recently determined a specific transcriptomic signature of chaperone-mediated autophagy in FAD astrocytes by using snRNA-seq of the FC of PSEN1 E280A FAD, SAD and healthy controls^18^. To determine whether astrocytic transcriptional states were consistent with the vascular proteomic alterations identified here, we re-analysed astrocytes from this snRNA-seq dataset. Eleven clusters were identified after the reanalysis of astrocytes from all groups. However, if every group is analysed independently, only five clusters can be identified in controls, nine clusters in SAD, and eight in FAD cases using the same parameters of clustering (Fig. 6A-B). Functional gene expression analyses of the five clusters identified in HC group showed pathways associated with astrocytic homeostatic functions such as cell junction organization, microtubule assembly and synaptic regulation, whereas other astrocytes showed a neuroinflammatory profile. Similar clusters can be also identified in SAD astrocytes, together with clusters associated with basement membrane, angiogenesis and circulatory system development. Notably, a cluster was identified associated with RNA regulation. Finally, FAD astrocytes also shared some functionally similar clusters either with HC controls or SAD subjects. Notably, astrocytic clusters functionally associated with RNA regulation and angiogenesis were present in both AD variants. FAD astrocytes showed unique clusters not apparent in the other groups, such as a cluster associated with glial cell differentiation, unfolded protein response, and ECM (Fig. 6C). These findings provide convergent transcriptomic support for altered RNA regulation in AD, as evident from our proteomic analysis, and the stress responses in FAD astrocytes^20^.

**Figure 6.**
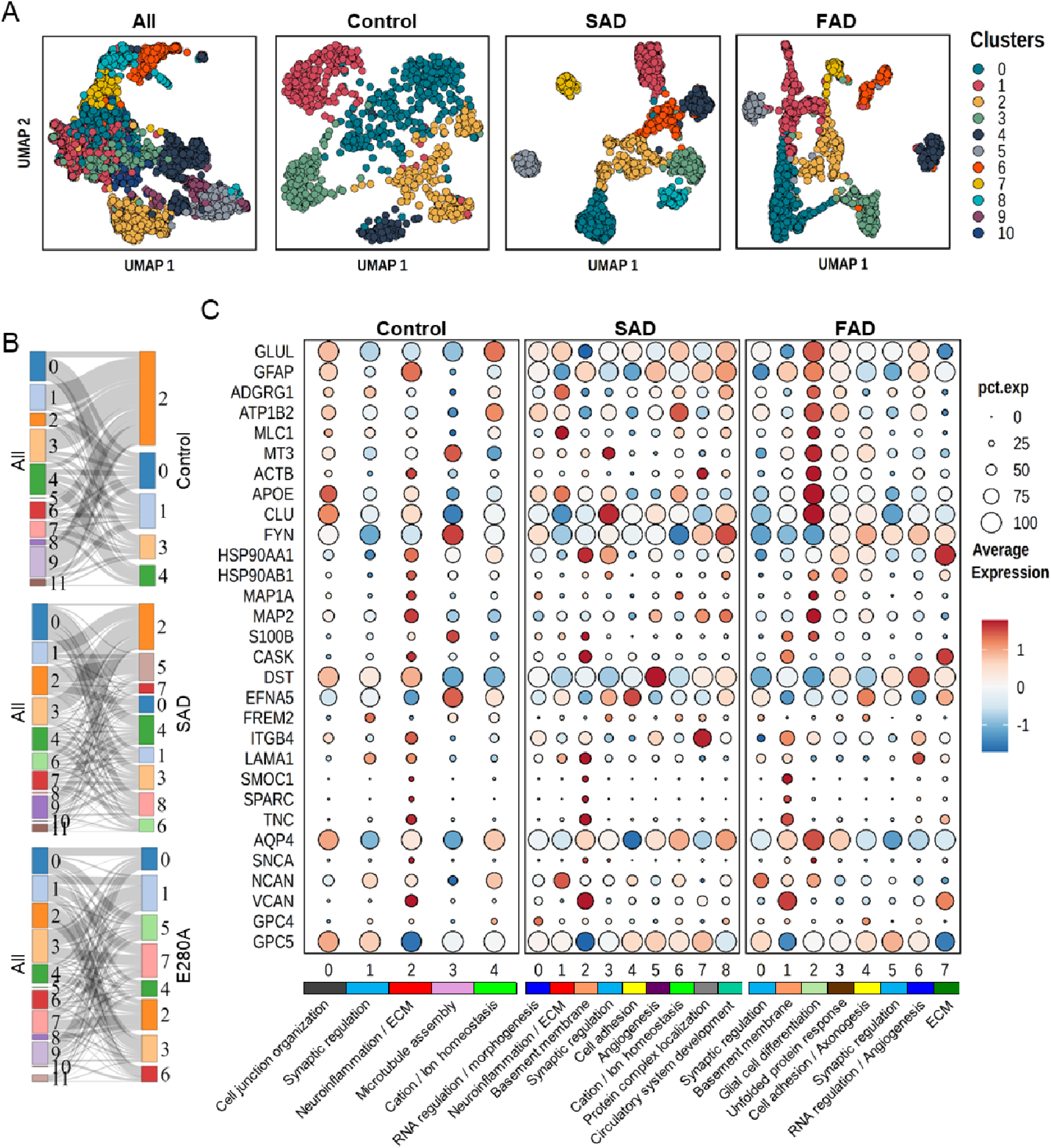
Abnormal single nuclei transcriptomic profile and subtypes of PSEN1 FAD astrocytes. A. Dimensionality reduction UMAP plots for all astrocytes studied in Almeida et al. (Ref 18), and the UMAP plots of the different groups (controls, n=8, SAD, n=8, and FAD, n=8) included in that study. Astrocytes were further clustered for our analysis. B. Sankey diagrams for the composition of the Control, SAD and FAD astrocytic clusters relative to the original clusters of all astrocytes studied. C. Dotplot heatmap according to percentage of expressing cells (pct. exp) and average expression for identifying genes and cluster composition for control, SAD and FAD astrocytes. Functional attributions for each cluster according to GSEA analysis and gene expression profiles are labelled under each cluster and group. Astrocytic clusters from both AD variants showed functional profiles for RNA regulation/morphogenesis, basement membrane, cell adhesion, and angiogenesis. FAD astrocytes additionally showed functional profiles in clusters for glial cell differentiation, unfolded protein response, RNA regulation, and extracellular membrane (ECM) genes.

## Discussion

Our central finding is that small vessel and perivascular astrocytic pathology in PSEN1 E280A FAD is an early event that is at least partly independent of Aβ and tau: a knock-in mouse lacking both pathologies reproduced the RNA-associated proteomic signature of human FAD microvessels and developed abnormal astrocytic end-feet by two months of age. We provide evidence for its multifactorial nature, with protein networks associated with features we have previously described in this population such as increased PVS^9^, CAA pathology^7^, or clinical features such as AoO or disease duration^10^. Similar previous studies identified known AD associated proteins, such as HTRA1, SMOC1, SMOC2, APOE, CLU, and COL1A1 dysregulated in MCI^28^, and CAA pathology with and without AD^27^. From a functional perspective both AD variants in our experiments also showed dysregulation of RNA-related proteins, and FAD also showed dysregulation of ECM, similarly reported in both MCI and AD^28^. It should be noted that by analysing independently proteins coprecipitating with Aβ aggregates in isolated microvessels, something that was not done in previous studies, we identified Aβ coprecipitating proteins to include mostly RNA biogenesis and RNA-related proteins complex assembly. Thus suggesting that some VSMCs already are expressing different RNA assemblies prior to Aβ deposition. Interestingly, even after removing Aβ coprecipitating proteins, RNA-related proteins remained as a prominent dysregulated feature in FAD, mostly in the OC, and correlating negatively with disease duration and CAA pathology severity. In the OC of FAD cases spliceosomal and ribosomal complex proteins were particularly affected. Furthermore, similar functional protein networks were dysregulated in a PSEN1E280A knock-in mouse model at relatively early age.

Ribosomal dysfunction and decreased protein synthesis as an early event of AD has been already described in AD brains^32^, and proteomic studies showed spliceosome components accumulation in insoluble fractions of AD and MCI brains^33^. More recently, cell cultures treated with Aβ 1-42 peptides showed downregulation of all spliceosome subunits^34^, and proteomic analyses in 6 months old APP/PS1 double transgenic mice showed dysregulation of ribosomal large subunit proteins^35^. These last two findings align with our identification of RNA-related proteins co-precipitating with Aβ aggregates in AD microvessels. In addition, small nuclear ribonucleoproteins specifically aggregate and colocalize with tau deposits in SAD and FAD cases^33^, subcellular fractionation of AD and control brains showed that ribosomes associate with tau in AD, leading to decrease of RNA translation^36^, and finally, spliceosomal components coimmunoprecipitates with tau in AD postmortem brains^37^. In summary, we provide strong evidence to suggest that the pathological accumulation of both, Aβ and tau, affects RNA metabolism and assembly in AD, and that it might be an early event in the pathogenesis. However, our PSENKi mouse model does not produce either of these protein aggregates, and yet it shows dysregulation of RNA-related proteins in isolated microvessels. Other transgenic mouse models including APP and tau mutations have shown common proteomic signatures with AD postmortem brains, particularly associated with the pathological hallmarks of the disease^38^. Recently, it has been shown in cellular and animal models that PSEN1 mutations induce U1 small nuclear RNA (snRNA) overexpression and loss of splicing function^39,40^. This provides a direct mechanistic route by which PSEN1 dysfunction, independent of amyloid, could produce the RNA-processing signature we observe in both human and murine microvessels. Our results support the impact of PSEN1 mutations in RNA-associated functions.

Besides RNA-associated proteins, ECM proteins showed to be specifically dysregulated in FAD microvessels and negatively correlated with PVS diameter in CAA affected vessels from those cases. ECM proteins have been found to be dysregulated in the parenchyma of brains from neurodegenerative diseases^41^, Down syndrome, and early and late AD^42^. ECM proteins upregulation is suggested to be also an early event in AD^43^, and to increase following disease progression according to Braak stages^44^. More to the point, ECM proteins were found to be dysregulated in MCI and AD microvessels^28^, and proteomics studies in astrocytes, a part of the neurovascular unit, have shown ECM proteins dysregulation^45,46^. These and our findings suggest that ECM dysregulation is another relevant feature in AD, and in SVD in AD. In fact, we also observed evidence of abnormal organization and decreased coverage of collagen in FAD microvessels. When vascular networks were assessed in FAD cases compared with healthy controls, increased lacunarity and presence of capillary kinking and knobbing were detected. Microvascular changes have been variously described in AD^47^, including abnormal angiogenesis^48^, evidenced by increased angiogenic markers in brain tissue^49^, and in CSF^50^. In SAD, abnormal angiogenesis can be attributed to multiple biological factors during the evolution of the disease^48^. In PSEN1 FAD, on the other hand, the underlying gamma secretase dysfunction can have an impact in angiogenesis, for instance by abnormal processing of NOTCH3, a known angiogenic factor^51^. Our morphological analyses in PSEN1Kimice confirmed the impact of this mutation in microvascular remodelling, evidenced by abnormal branching and tortuosity. Thus, we consider abnormal angiogenesis to be a main SVD feature in PSEN1 FAD, supported not only by our morphological findings but also by its proteomic signature, given that ECM plays a key role as a modulating factor in vascular remodelling^52^, astrocyte-derive remodelling factors have shown to be dysregulated in AD^53^, and RNA-binding proteins and RNA translation are modulators of angiogenesis^54^.

Astrocytes play critical roles in synaptic, metabolic and structural support of different functional systems in the central nervous system. Among them, the neuro-glia-vascular unit is a structural and functional regulator of blood flow and the basic unit of the Blood Brain Barrier (BBB). Astrocytes modulate BBB permeability and vascular flow by the release of vascular factors and spatial connections with endothelial cells via their end-feet^55^. In neurodegeneration, reactive astrocytes can turn pro-inflammatory or anti-inflammatory. In AD, both types of astrocytes are associated with different disease features, including synaptic dysfunction, amyloid plaque clearance and neurovascular dysfunction^56,57^. We recently detected involvement of astrocytic dysfunction in vascular pathology in PSEN1 E280A^9,31^, their single cell transcriptomic signature^20^, and a main role in the mechanisms of protection generated by the APOE Christchurch mutation^30^. In fact, one of the notable features of the homozygous APOE Christchurch carrier affected by the PSEN1 E280A mutation was their uncharacteristically sparse CAA pathology^30^, a feature shared partially by APOE Christchurch heterozygotes from the same family^58^. So far these findings, either pathological or protective, were considered as directly related to AD known mechanisms, such as Aβ or tau aggregation. In the current study we obtained evidence, for the first time, of perivascular astrocytic pathology as the earliest event in a murine model for PSEN1 E280A, in absence of any other AD pathological feature. This, together with a new analysis of the astrocytic snRNAseq data obtained from these patients, and the proteomic signature of PSEN1 E280A cerebral microvessels, suggest perivascular astrocyte pathology and SVD as early events in PSEN1 E280A FAD pathology.

This study supports a multifactorial model for the inherent cSVD pathology in AD. Most previous studies focused on this question examined end of stage disease. Therefore, several years of pathology, including the accumulation of Aβ peptides in vessel walls, together with all the different molecular events involved in inflammation and neurodegeneration, contribute to the pathology evidenced in the proteomic signature of this disorder. However, we found additional proteins and pathways affected in FAD cases, a distinct proteomic signature in their FC, and higher number of dysregulated proteins in the OC. We found that RNA processing and translation are commonly dysregulated processes among both AD variants and our FAD knock-in model. Our morphological analyses suggest that these pathways are related to abnormal angiogenesis in PSEN1 mutants. Thus, several pathological events, either CAA pathology or abnormal PSEN1 activity, could contribute to abnormal angiogenesis in FAD. In addition, the PSEN1 E280A mutation presented with early astrocytic damage in PSEN1Ki mice, and we identified human astrocytic single cell clusters involving similar biological processes than those detected in our proteomic analyses, therefore confirming another layer of pathological events associated with SVD in FAD.

Limitations of our study include the intrinsic difficulty of studying molecular mechanisms of disease in postmortem samples of end of stage pathology in AD. We attempted to reduce the uncertainty by analysing PSEN1Ki mice lacking APP or tau associated pathologies. Furthermore, variability of AD pathology presentation among individual cases can affect our results and complicate their interpretation. Further analyses in a larger number of cases or with a more uniform disease presentation might improve the accuracy of such studies. The development of more precise cellular or animal models recapitulating SVD in AD would clarify further the involvement of each molecular pathway identified.

The relevance of our findings for the pathological progression and severity of FAD is further highlited by the protection against brain vascular disease identified in the PSEN1 E280A homozygous Christchurch case^30^, and partially shown by PSEN1 E280A heterozygous Christchurch cases^58^, given that the regions in which less vascular pathology was identified were the most protected against tau pathology and neurodegeneration. It has already been suggested that SVD is an early event in AD^8^. For instance, lower blood flow has been identified in prodromal AD which may lead to selective chronic hypoperfusion ^59,60^, and cognitive decline^61^. PSEN1 E280A FAD cases show a prodromal stage of disease around 15 years before dementia onset^10^. However, PSEN1 E280A carriers in their second decade of life already show increased levels of Aβ 1-42 in plasma as an early phenotype of the pathology^62^. In our PSEN1 E280A knock-in mouse model, in absence of human APP, abnormal perivascular astrocytic end-feet were the first phenotypic trait we detected, later to be associated with increased vascular wall thickness. Given that loss of BBB integrity has been associated with higher levels of Aβ 1-42 in plasma^63^, it could be suggested that Aβ 1-42 plasma levels in young PSEN1 E280A carriers can also be indirectly associated with brain microvascular damage. Further studies need to address this issue and determine further the early role of SVD in FAD. Finally, our study shows that several factors and biological protein networks play a role in SVD in all AD. This concept reinforces the necessity of early intervention for cardiovascular risk factors in middle aged individuals, and the incorporation of therapeutic approaches aimed at SVD in early stages of dementia.

## Acknowledgements

The authors would like to acknowledge the invaluable contribution of the Colombian families suffering from Familial Alzheimer’s disease for donating biological samples for this study. We thank the staff and technical support of the UKE imaging facility (UMIF), and the Core Facility Mass Spectrometric Proteomics at UKE. This work was funded by the Deutsche Forschungsgemeinschaft (DFG) with project number 458854216 to D.S-F. By the NINDS and NIA cofunded grant no. RF1 NS110048 to J.A.-V. and D. S.-F., and by a gift Coefficient Giving and Good Ventures to J.A.-V., L.L.K., S.K., and D. S.-F. RK’s work was supported by grants from the UK Medical Research Council (MRC, G0500247), Newcastle Centre for Brain Ageing and Vitality (BBSRC, EPSRC, ESRC and MRC, LLHW), and Alzheimer’s Research UK (ARUK).

## Contributions

D. S.-F. and J.A.-V. initiated this work, D. S.-F., J.A.-V., R.K., M.E., L.L.K., R.P.-D. and F.L. supervised it. J.L.L., K.-Y.K., N.D.V.-M., C.O.-C., S.W., J.A.O., I.S.-U. and S.K. performed experiments and collected data. J.L.L., K.-Y.K., J.A.O., I.S.-U., R.P.-D., and D.S.-F. performed general data analysis. Y.E.H.E.A., M.S., C.O.-C., S.W., and D.S.-F. performed proteomic and transcriptomic analysis. D.C.-M., A.V., M.G., and F.L. performed clinical and pathological diagnosis. J.L.L., Y.E.H.E.A., K.-Y.K., R.P.-D., M.E., and D.S.-F. drafted the manuscript. All authors corrected it and approved it. J.L.L. and Y.E.H.E.A. are equal contributors listed as first coauthors.

## Abbreviations

AD: Alzheimer’s disease
Aβ: Amyloid-beta
APP: Amyloid precursor protein
BBB: Blood Brain Barrier
CAA: Cerebral amyloid angiopathy
cSVD: Cerebral Small Vessel Disease
ECM: Extracellular Matrix
FAD: Familial Alzheimer’s disease
PVS: Perivascular Space
SAD: Sporadic Alzheimer’s disease
SVD: Small Vessel Disease
NFTs: Tau neurofibrillary tangles
WGCNA: Weighted Gene Correlation Network Analysis

