## Supplementary material for "Astrocyte-driven small vessel disease is an early, amyloid-independent feature of PSEN1 E280A familial Alzheimer’s disease"

### Villalba-Moreno et al. Supplementary Files

#### Supplementary Figure legends

**Supp. Fig. 1.** Generation and genotyping of the PSEN1 E280A knock-in mouse model (A) Sequence design for the sgRNA and resulting sequence modifications in transgenic animals. (B) Genotyping of a homozygous PSEN1 E280A knock-in mouse.

**Supp. Fig. 2.** Peptidomic analysis of Ab peptides. Aβ abundances are shown for all the comparisons. Comparisons between Familial Alzheimer's Disease (FAD) (n = 10) and Sporadic Alzheimer's Disease (SAD) (n = 5). In the Frontal Cortex (FC) Aβ37 and Aβ38 were significantly higher in FAD group, whereas Aβ42 and Aβ40 were not different. Aβ42/Aβ40 ratio was found significantly different between groups (A). Comparisons between FAD (n = 10) and SAD (n = 5) in Occipital Cortex (OC), and Aβ42/Aβ40 ratio between these groups showed no significant differences (B). Comparison between FAD in FC versus OC showed no differences, whereas Aβ42/Aβ40 ratio was found to be significantly different between the groups (C). Comparison between SAD in FC versus OC, and Aβ42/Aβ40 ratio between these groups showed no significant differences (D).

**Supp. Fig. 3.** Changes in proteoglycans and heparin-binding proteins in FAD. (A) Venn diagram showing the overlap between dysregulated proteins in FAD occipital (O_FAD_vs_O_HC), FAD frontal (F_FAD_vs_F_HC), and proteoglycans. (B-C) Volcano plots depict the fold change of each protein compared to the control for (B) FAD occipital and (C) FAD frontal regions. Proteoglycans are highlighted in red, and select proteins described as interacting with proteoglycans are depicted in blue. (D) Gene Ontology classification of dysregulated proteins in FAD occipital region based on molecular function was performed using DAVID web resource. The histogram reports on number of genes and -log (p-value). (E) Scatter plot of FAD occipital region dysregulated genes classified as heparan sulfate-binding in (D).

**Supp. Fig. 4.** Enrichment analysis from DEPs. Biological Processes (BP) of Gene Ontology (GO) database resulting from the list of common differential expressed proteins (DEPs) in Familial Alzheimer's Disease (FAD) and Sporadic Alzheimer's Disease (SAD) versus Controls in Frontal Cortex (FC) with at least 2 common genes are shown as a bar plot and a heatmap (A). Barplots of the top 8 GO terms of each category obtained from the common DEPs in FAD and SAD versus Controls in FC (B). Barplots of the top 8 GO terms of each category obtained from the unique DEPs in FAD versus Controls in FC (C). Barplots of the top 8 GO terms of each category obtained from the unique DEPs in SAD versus HC in FC (D).

**Supp. Fig. 5.** Module 7 of WCGNA in FAD cases. Functional enrichment analysis of module eigengenes 7 (ME&) that significantly correlated in WGCNA of FAD cases. The results are shown as a barplots of the top 8 terms of each Gene Ontology (GO) category.

**Supp. Fig. 6.** Distribution of SMOC1 in AD microvessels. (A) Representative images are shown for 50 um caliber arterioles in the FC and OC in FAD and SAD (n=3). Sections were stained for DAPI, A□, SMOC1 and GFAP, scale bar = 50μm. In FC FAD, there was SMOC1 in the vessel wall. In the OC of FAD SMOC1 staining could be observed within the vessel wall and in the surrounding plaque. Similarly, in the FC of SAD

SMOC1 signal is restricted to the vessel wall. In the OC of SAD SMOC1 signal was observed in the vessel wall and in the plaque. (B) There were no differences in SMOC1 abundance in the proteomic analysis between FAD and SAD.

**Supp. Fig. 7.** Distribution of Tenascin C in AD microvessels. (A) Representative images for DAPI, aquaporin-4 (AQP4), tenascin C (TNC) and collagen IV were stained in FAD and SAD FC and OC sections (n=1), scale bar = 50µm. In FAD the FC and OC did not present differently in the TNC staining. In the FC of SAD, TNC was localised in astrocytes (white arrowheads). In the OC of SAD TNC distributed everywhere as in FAD. (B) The abundance levels of TNC in the proteomics of purified vessels did not differ between FAD and SAD but the FC of SAD had the highest abundance.

**Supp. Fig. 8.** Cell type deconvolution of studied proteins in AD microvessels. Heatmap of cell type deconvolution analysis showing the relative estimation scores for each case and cell type of interest (astrocytes, endothelial cells, pericytes, smooth muscle cells). Each case is further annotated based on age of death, sex and condition.

**Supp. Fig. 9.** Vascular morphology analysis of WT and PSEN1 E280A Knock in mice. (A) Representative images are shown for laminin-stained paraffin-fixed sections for 24-week-old WT, (B) 24-week-old PS1KI and (C) 52-week-old PS1KI (n=3). (D) Similarly, representative images for SMA are displayed for 24-week-old WT, (E) 24-week-old PS1KI and (F) 52-week-old PS1KI, scale bar = 100µm. (G) The %area of laminin and (I) SMA was quantified, and no significant difference was found. (H) Further, there was no significant difference in the integrated density of laminin or (J) SMA. (K) However, in 24-week old PS1KI there were significant more tortuous vessels compared to 24-week-old WT and 52-week-old PS1KI (p: ****<0.001, **=0.0022).

**Supplementary Figures**

#### Supplemental Figure 1

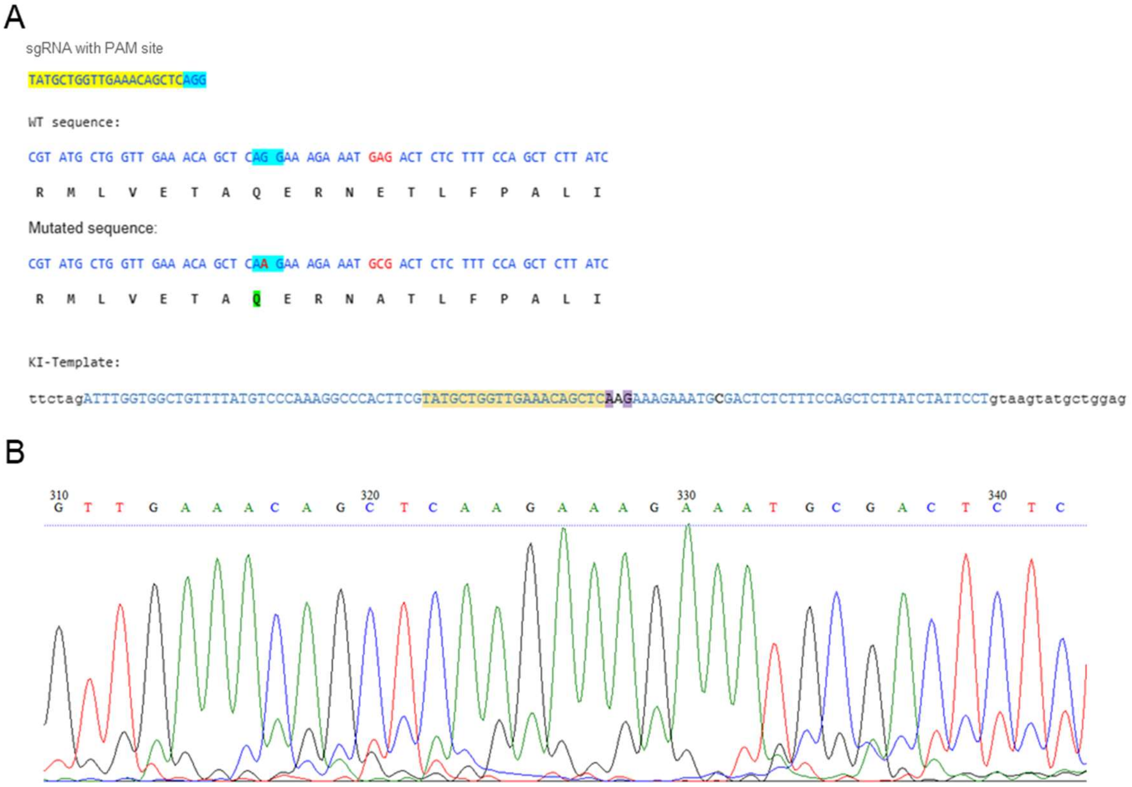

**Supplemental Figure 2**

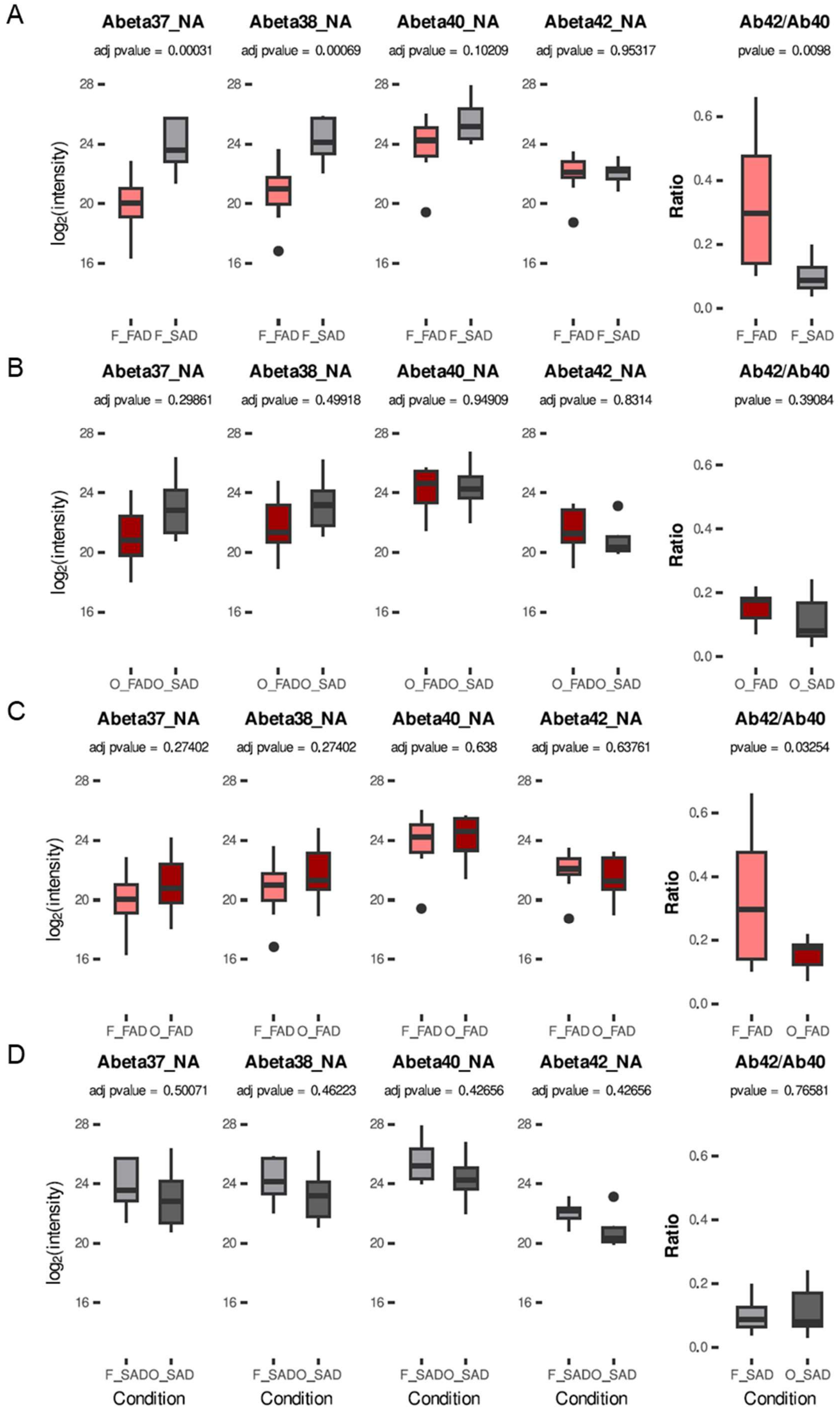

#### Supplemental Figure 3

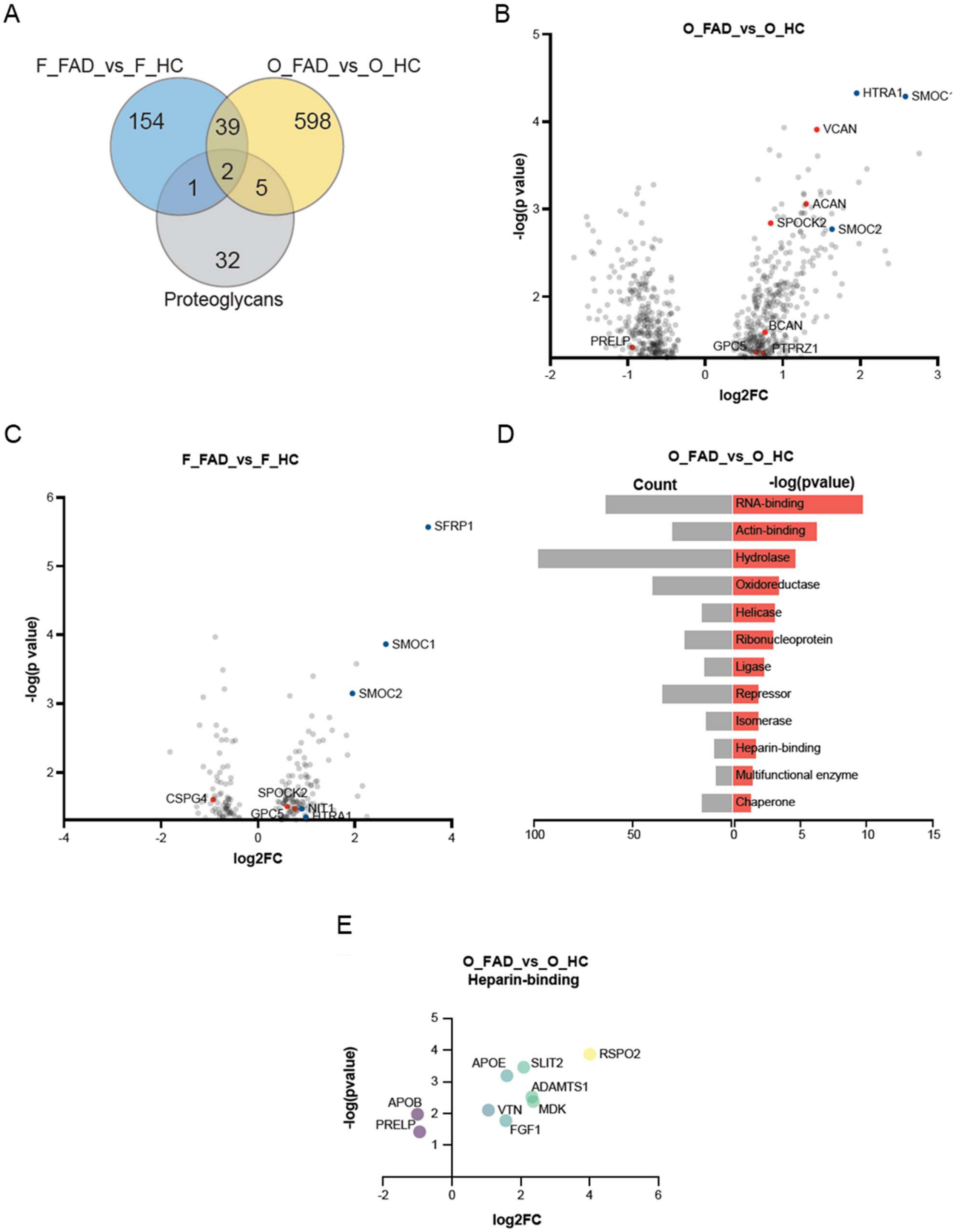

#### Supplemental Figure 4

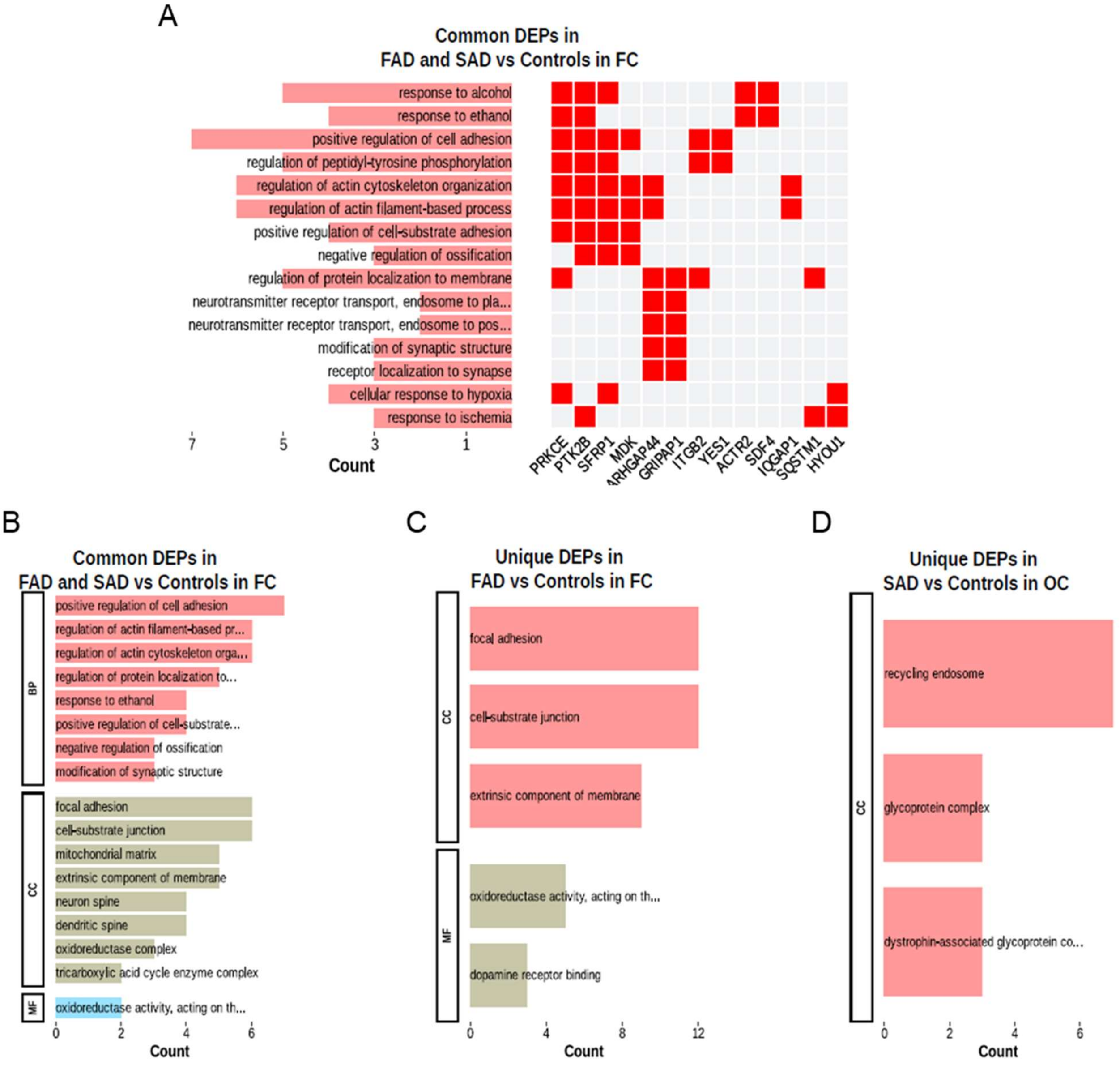

**Supplemental Figure 5**

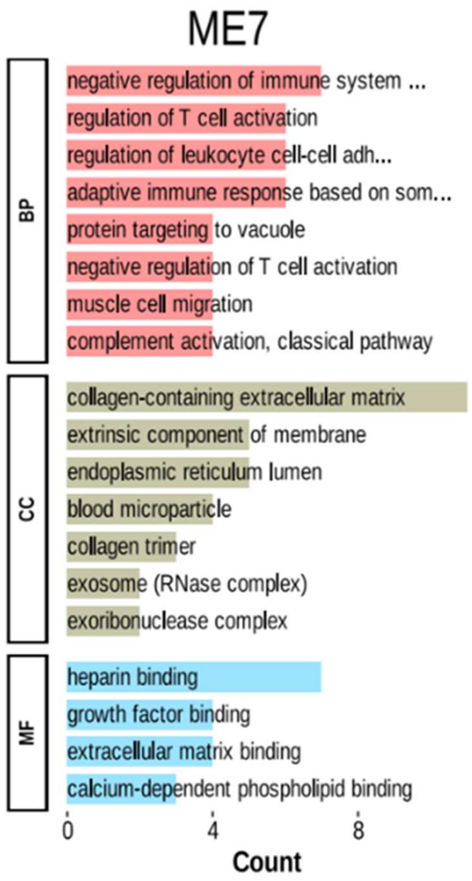

#### Supplemental Figure 6

A

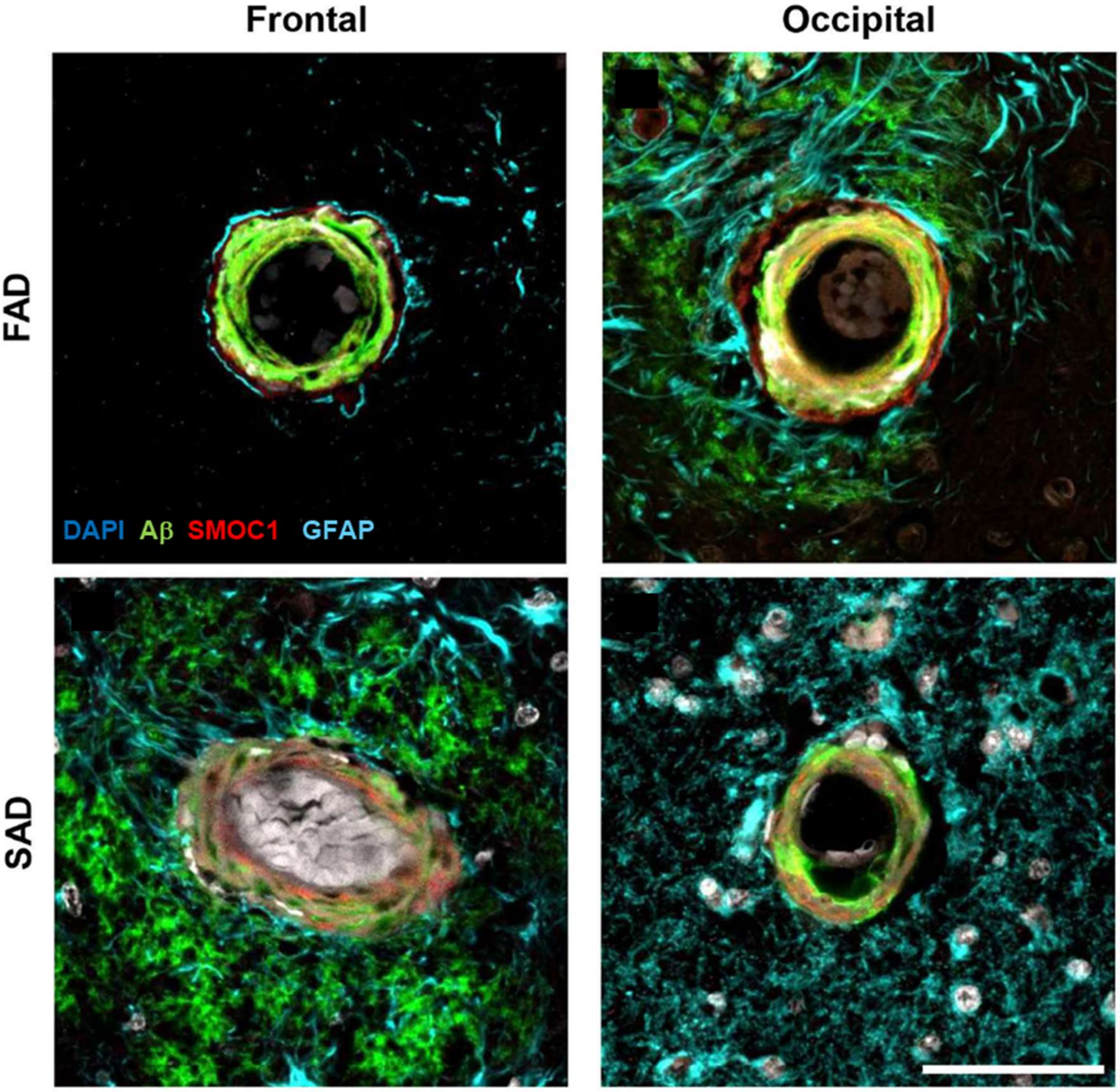


B

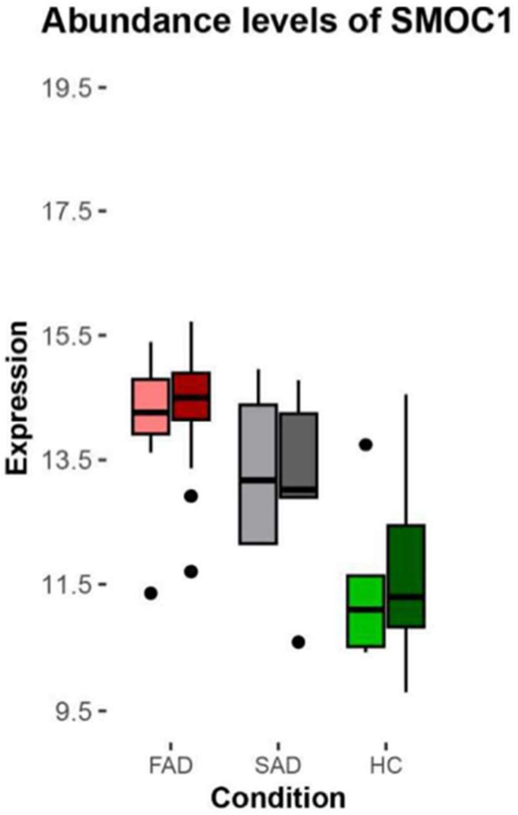

#### Supplemental Figure 7

A

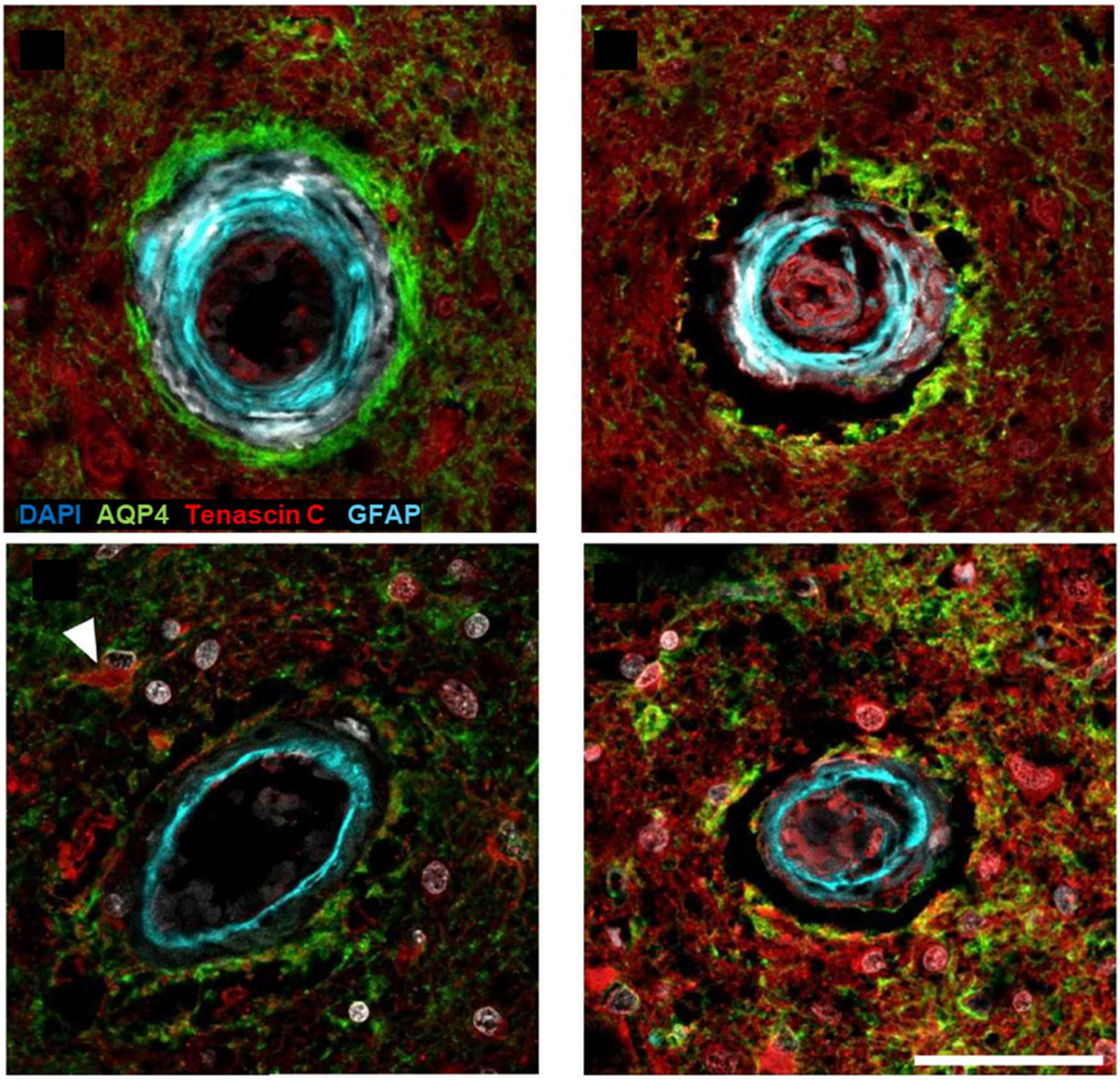


B

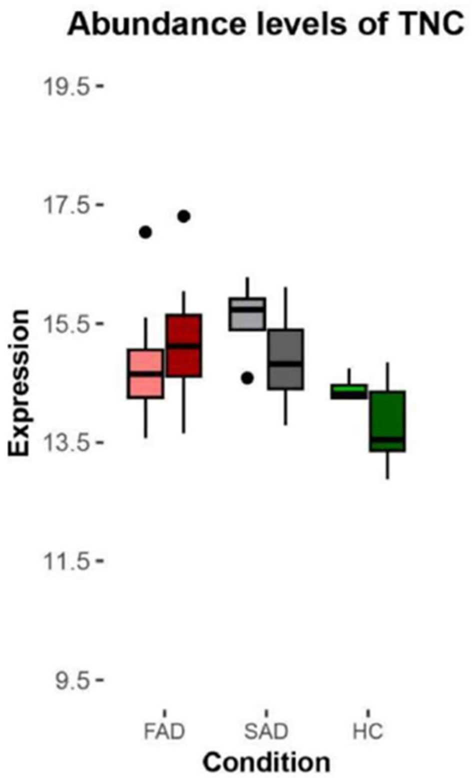

#### Supplemental Figure 8

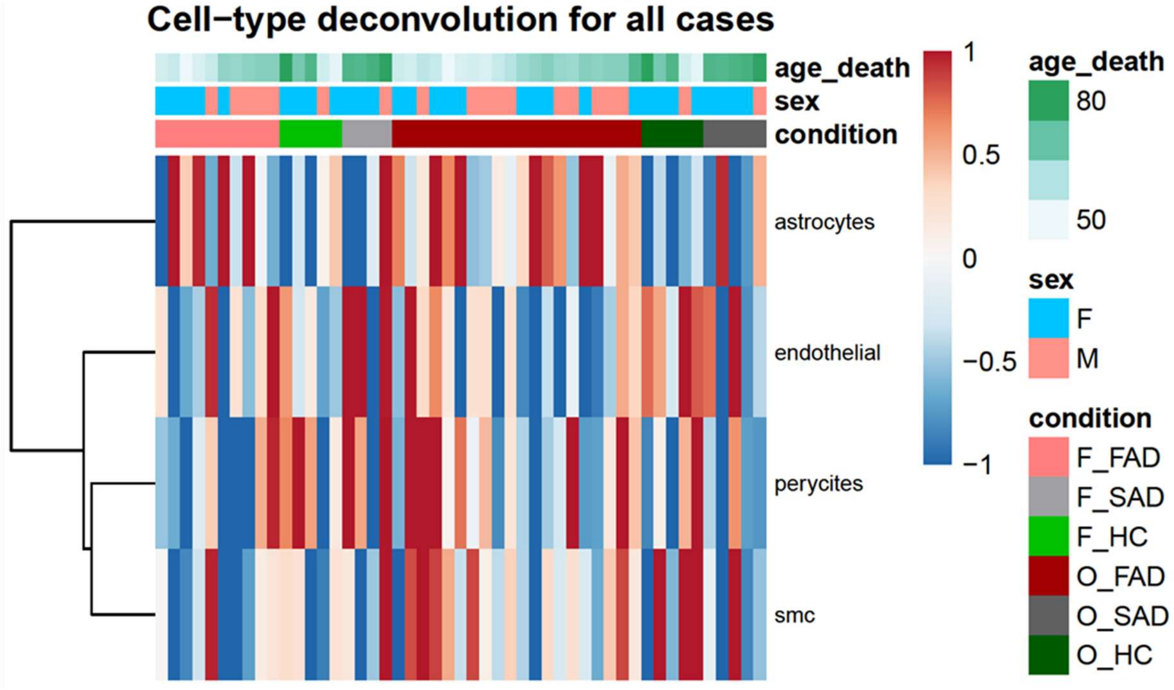

### Supplemental Figure 9

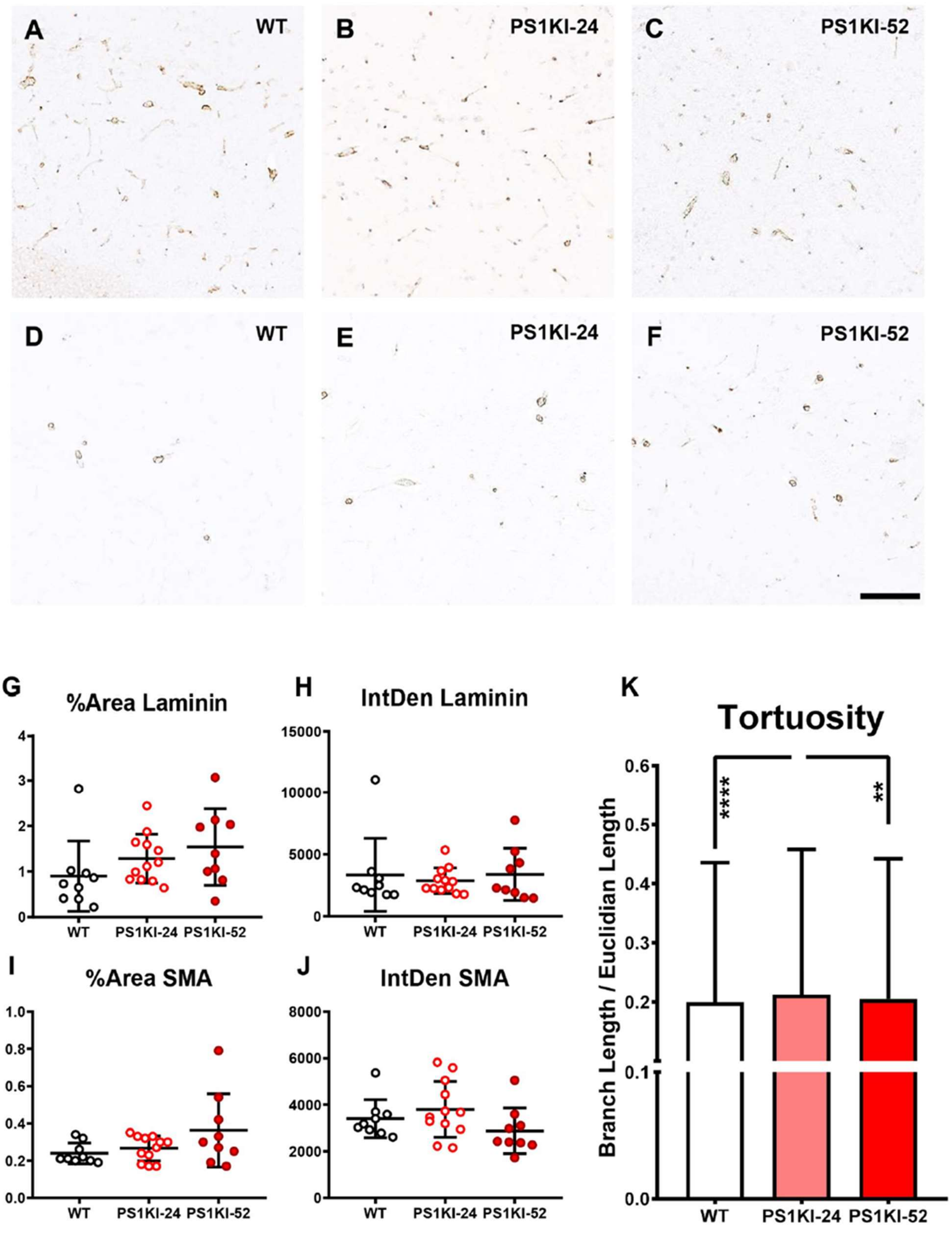
